# ASTK: a machine learning-based integrative software for alternative splicing analysis

**DOI:** 10.1101/2023.01.03.522470

**Authors:** Shenghui Huang, Jiangshuang He, Lei Yu, Jun Guo, Shangying Jiang, Zhaoxia Sun, Linghui Cheng, Xing Chen, Xiang Ji, Yi Zhang

## Abstract

Alternative splicing (AS) is a fundamental mechanism that regulates gene expression. Splicing dynamics is involved in both physiological and pathological processes. In this paper, we introduce ASTK, a software package covering upstream and downstream analysis of AS. Initially, ASTK offers a module to perform enrichment analysis at both the gene- and exon-level to incorporate various impacts by different spliced events on a single gene. We further cluster AS genes and alternative exons into three groups based on spliced exon sizes (micro-, mid-, and macro-), which are preferentially associated with distinct biological pathways. A major challenge in the field has been decoding the regulatory codes of splicing. ASTK adeptly extracts both sequence features and epigenetic marks associated with AS events. Through the application of machine learning algorithms, we identified pivotal features influencing the inclusion levels of most AS types. Notably, the splice site strength is a primary determinant for the inclusion levels in alternative 3’/5’ splice sites (A3/A5). For the alternative first exon (AF) and skipping exon (SE) classes, a combination of sequence and epigenetic features collaboratively dictate exon inclusion/exclusion. Our findings underscore ASTK’s capability to enhance the functional understanding of AS events and shed light on the intricacies of splicing regulation.

## Introduction

Alternative splicing (AS) is an essential mechanism to increase transcriptomic and proteomic diversity. In humans, over 95% of multi-exonic genes undergo this process, producing a myriad of splice isoforms from a single mRNA [1]. While alternative splicing is critical for normal physiological functions, aberrations in this process can lead to a spectrum of diseases [2]. Numerous software tools have been crafted to aid AS research, ranging from the detection and quantification of AS events [3–9], to time series and isoform switch analysis [10, 11], functional enrichment [12, 13], mapping the spatial distribution of RBPs (RNA binding proteins) binding motifs [14], sequence feature extractions [15, 16], and assessing the functional implications of AS changes [16]. Historically, AS studies have been focusing on two main aspects: functionality and regulation. However, researchers usually studied the two aspects separately, due to the lack of comprehensive tools. Studying both aspects together will advance our understanding of mis-splicing events and help us design novel therapeutic approaches. For this purpose, by employing the most up-to-date algorithms and techniques, we developed a comprehensive software package to address the needs: i) characterizing the distribution patterns of AS events, ii) systematically analyzing functional effects of AS events at the gene- and exon-level respectively, iii) dissecting out splicing-associated sequence features and epigenetic marks, and iv) inferring the splicing regulatory code for each type of AS.

AS plays a vital role in biological functions, influencing cellular processes, tissue specificity, developmental stages, and disease conditions [1, 17]. Traditionally, AS analysis is centered on genes. It is noteworthy that this gene-central approach may bring many potential pitfalls since not all spliced exons have a functional impact. In addition, a single gene can have multiple AS events simultaneously, which may affect the gene’s functions differently. Therefore, we also need to investigate the AS events at the exon level. In ASTK, we provide a module integrating the network-based enrichment method (NEASE) for AS events to perform the functional enrichment analysis of AS at both the gene- and exon-level [12].

Under evolutionary constraints, splicing is tightly regulated to ensure correct forms of the pre-mRNAs. Exon conservation, both in sequence and length, has been implicated in this regulation [18]. Statistically, the majority of spliced exons in human and mouse genomes are less than 300 nucleotides (nt) [19, 20]. Comparative genomics of 76 vertebrate species, including humans, demonstrated that exons of less than 250 nt are highly conserved in length. And more importantly, there exists a strong correlation between exon length and exon sequence conservation for these exons [18], suggesting that exon size is an evolutionarily conserved feature. Meanwhile, exons longer than 250 nt are low in length conservation [18]. Another study identified over 13,000 micro-exons (≤ 51 nt) in the human genome, which are highly conserved across species [21]. These micro-exons have been linked to tissue-specific expression and are pivotal in processes such as cell differentiation, migration, and neural functions [22–24]. Dysregulation of micro-exons has been associated with neurological disorders and cancers [25]. In our research, we categorized AS events based on spliced exon lengths: micro-exons (≤ 51 nt), mid-sized exons (52-250 nt), and macro-exons (≥ 251 nt). At the gene level, after clustering, we found that the three groups are likely enriched in distinct biological pathways in our case studies. Similar observations were made at the exon level too. Such trends become clearer when AS events classified into individual types.

The splicing of pre-mRNA is orchestrated by the spliceosome, a complex comprising five core snRNPs and over 300 different RBPs [26]. As a highly regulated process, the decision as to which exon is included or removed depends on intricate interactions among RNA sequence elements, splicing regulators, and epigenetic components. Sequence features like splice site strength, exon/intron architecture, and GC contents are critical for exon recognition and selection [27, 28]. Additionally, factors such as histone modifications, chromatin conformation, and accessibility influence splicing outcomes [1, 29–35]. Although an increasing number of studies have established the link between sequence and epigenetic features and AS, these studies are typically restricted to test the association of skipping exon (SE) with one type of the features at a time. Besides SE and retained intron (RI) [34, 36], the regulatory mechanisms in other types of AS are largely unexplored. Just as important, how sequence and epigenetic features work together to control divergent AS remains elusive. With ASTK, we systematically investigated the sequence features and epigenetic marks associated with each AS class in our test cases. Intriguingly, we uncovered key features that significantly impact exon inclusion levels across six classes except for mutually exclusive exons (MX): alternative 3’ splice site (A3), alternative 5’ splice site (A5), alternative first exons (AF), alternative last exons (AL), RI, and SE. Moreover, the influence of sequence and epigenetic markers varied substantially across AS classes, suggesting the existence of distinct regulatory mechanisms for different types of AS events.

With the development of high-throughput sequencing technologies and the reduction of cost, huge amounts of transcriptomic data accumulated up to now have provided unprecedented resources and challenges to fully characterize AS events. Despite multiple software packages available for AS analysis, most of them focus on only one or few aspects. Here, we present a new software package, ASTK, with a command-line interface for comprehensive analysis of AS, including clustering AS events based on the size of spliced exons, differential AS analysis, functional enrichment analysis at the gene- and exon-level separately, extracting sequence and epigenetic features, identifying RBP binding motifs, inferring splicing effects of RBPs, and machine learning-based modeling for feature ranking and splicing code decryption. We demonstrated the use of ASTK with published multi-omics data. Our results indicate that results of function enrichment analyses of AS events are different and can be complementary at the gene- and exon-level. In terms of AS regulation, sequence features emerged as primary influencers for most AS classes during embryonic mouse forebrain development. Specifically, splice site strength is a key determinant for A3 and A5 exon inclusion levels. Additionally, beyond sequence features, specific histone modifications and open chromatin states play pivotal roles in exon selection and usage for AF and SE. Therefore, given the dynamic nature of transcription regulation, ASTK will aid in characterizing the splicing code in a spatial-temporal, condition-specific manner.

## Materials and methods

The components and workflow of ASTK are graphically depicted in Figure 1A. Comprising multiple modules, ASTK is designed for user-friendly operation. Detailed instructions for each module are readily available in the online manual. Each function within these modules is scripted to offer a command-line interface, allowing users to effortlessly initiate and tailor the scripts to their preferences.

**Figure 1.**
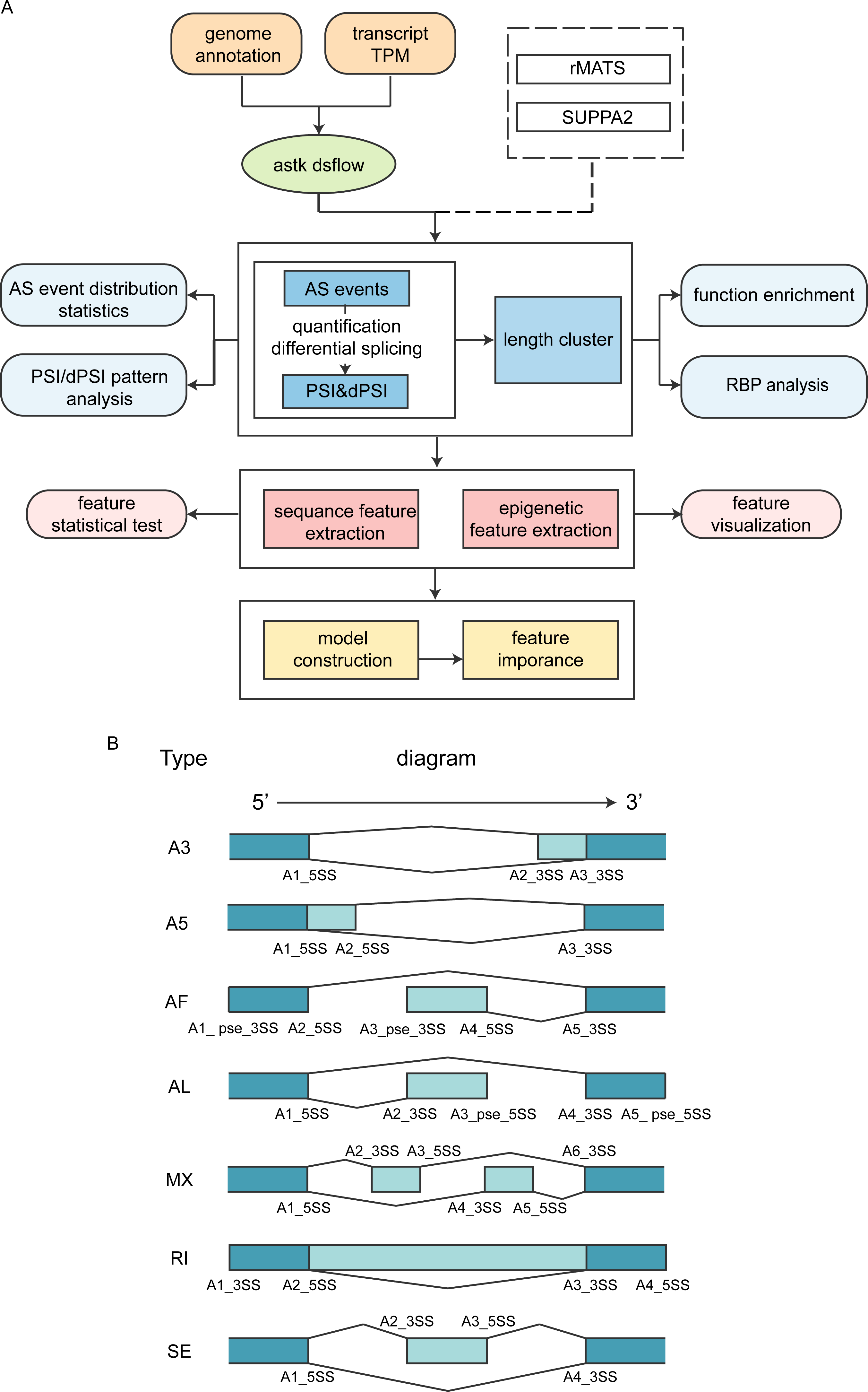
ASTK workflow. **A.** Schematic illustration of the workflow **B.** Diagram of seven AS types: alternative 3’ splice site (A3), alternative 5’ splice site (A5), alternative first exons (AF), alternative last exons (AL), mutually exclusive exons (MX), retained intron (RI), and skipping exon (SE). “A1” “A2”, “A3”, “A4”, and “A5” denote the order of splice sites in AS events. “5SS” and “3SS” denote the 5’ and 3’ splice site, respectively.

### AS events detection, quantification, and differential splicing analysis

Among the myriad of AS tools available, rMATS and SUPPA2 have consistently emerged as top choices. Recent benchmarking, which compared 21 differential splicing tools, ranked SUPPA2 and rMATS among the best performers [37]. While conventional tools like rMATS classify AS events into five distinct types: A3, A5, MX, RI, and SE, SUPPA2 introduces two additional classifications: AF and AL, bringing the total to seven. Given this expanded categorization, we opted for SUPPA2 for our preliminary AS analysis.

As illustrated in Figure 1A, we developed built-in functions based on SUPPA2 [4] for AS events detection and quantification, with flexibility for samples under different conditions. Also, ASTK can utilize output files generated by other programs, including SUPPA2 and rMATS [3], for downstream analysis. According to the program guide, differential AS events can be selected using cutoffs of ΔPSI (Percent Spliced-In), p-values, or FDR values. Furthermore, users have the option to cluster AS genes and spliced exons by the lengths of alternative exons. For data interpretation, the top 25% and bottom 25% based on PSI values are designated as PSI high and PSI low, respectively. Users can customize these thresholds, opting for other delineations such as the top 33% and bottom 33%, or the top 40% and bottom 40%.

### AS events distribution pattern analysis

ASTK incorporates a suite of functions designed to compute statistics and analyze the expression patterns of AS events. Within the software:

- **Barplots** are employed to represent the frequencies of various AS event types across different experimental conditions.
- **UpSet plots** offer a visual representation of the shared and unique AS events between conditions.
- **Principal Component Analysis (PCA)**, utilizing percent spliced-in (PSI) values, enables visualization of sample variations and serves as an effective tool for quality control.
- **Heatmaps** of PSI values facilitate rapid identification of notable changes in AS events between samples.
- **Volcano plots**, plotting ΔPSI values on the x-axis and −log10(p-values) on the y-axis, provide insights into statistically significant AS events.

### Functional enrichment analysis for AS events

Within ASTK, we’ve incorporated three functional enrichment methodologies: Over-Representation Analysis (ORA), Gene Set Enrichment Analysis (GSEA), and the Network-Based Enrichment Method for AS Events (NEASE) [12].

- **ORA and GSEA**: These methods facilitate gene-level enrichment analysis for genes exhibiting differential alternative splicing. This analysis is executed utilizing the clusterProfiler R package [38]. Notably, ORA can sometimes yield redundant Gene Ontology (GO) terms. To streamline these results, we’ve integrated the simplifyEnrichment R package [39], ensuring more concise and meaningful enrichment outcomes.
- **NEASE**: This method is tailored for exon-level enrichment analysis. It’s adept at pinpointing the structural consequences of spliced exons and elucidating the subsequent remodeling of protein-protein interactions [12].

### RBPs motif analysis and splicing effects

ASTK is equipped to carry out RBP binding motif enrichment analysis, focusing on the flanking regions adjacent to 5’ and 3’ splice sites. To achieve this, the software employs the CentriMo algorithm [40], leveraging the CISBP-RNA and ATtRACT motif databases [41, 42]. Identified motifs of significance are visually represented as sequence logos, with the accompanying density curve depicting the probability of the given motif at each position. Furthermore, ASTK incorporates a feature that allows users to summarize and discern pattern shifts across various AS event types under specific conditions, such as RBP knockdowns or overexpression.

### Sequence and epigenetic feature extraction

In ASTK, we’ve integrated a suite of functions designed to extract both sequence and epigenetic features pertinent to AS events. At present, ASTK is capable of computing attributes such as exon/intron lengths, GC contents, and splice site strength across all AS types. Users have the flexibility to determine the flanking regions of each splice site for which GC content is calculated.

Building upon the MaxEntScan [27] Perl script, we’ve crafted a function in ASTK that computes the strength of splice sites for any given AS event. The underlying maximum entropy model takes into account both adjacent and non-adjacent positional dependencies. Strength scores are derived using a log-odds ratio, applicable to either a 9-mer sequence (for 5’ splice sites) or a 23-mer sequence (for 3’ splice sites) [27].

On the epigenetic front, ASTK adeptly extracts features, encompassing aspects like histone modifications and chromatin accessibility. To correlate specific epigenetic features with AS events, ASTK calculates read coverage along the regions flanking the 5’ and 3’ splice sites. This is achieved by using mapped reads from each ChIPseq or ATACseq sample. The read alignment BAM files undergo normalization and are subsequently converted into the bigwig file format, with the user’s preferred bin size.

### Feature visualization and statistical comparison

ASTK provides a function designed specifically for the statistical comparison of individual sequence features across conditions, leveraging multiple statistical test methods. The software vividly displays the distribution of epigenetic signals in regions flanking splice sites, using heatmaps centered on each splice site.

### Machine learning modeling and feature importance

ASTK introduces a machine learning module, built upon the Scikit-learn API [43], which supports the random forest classifier. To optimize model performance, ASTK offers the option of grid search for hyperparameter tuning. The software employs cross-validation to assess the efficacy of the models. Various classical metrics, including accuracy, AUC, recall, precision, and F1 scores, are computed within ASTK to gauge model performance.

For enhanced model interpretability, feature importance is quantified using the Gini importance measure. This metric tallies the reduction in impurity each feature induces across all trees in the model — a higher value indicates greater feature importance.

The formula to compute the Gini importance for a feature in a Random Forest model is: *Gini importance (k) = Σ (Split node count / Total node count) * (Gini impurity (parent) - Gini impurity (child))*

Where:

- *Split node count*: Refers to the count of nodes where feature *k* is utilized for splitting.
- *Total node count*: Denotes the overall node count within the Random Forest.
- *Gini impurity (parent)*: Represents the impurity of the node pre-split.
- *Gini impurity (child)*: Indicates the impurity of each descendant node post-split.

### Availability of software and data

ASTK is crafted in Python and is readily accessible on the Python Package Index (PyPi), Python’s official third-party software repository. A simple installation can be achieved using commands such as pip (available at: https://pypi.org/project/astk/). For those seeking ease of configuration and minimized dependency issues, ASTK is also encapsulated within a Docker image, freely obtainable from DockerHub at https://hub.docker.com/r/huangshing/astk. Comprehensive online guidance is provided at https://huang-sh.github.io/astk-doc/. The datasets used in this study, including their descriptions and accession numbers, are cataloged in Supplemental Table 3.

## Results

### Testing ASTK with published datasets

In this section, we used public data to evaluate the performance of ASTK, including multi-omics datasets of C57BL/6 mouse forebrain development (ranging from E11.5-P0) downloaded from ENCODE, RNAseq datasets of human embryonic stem cells at seven early neural differentiation time points (from 0 to 72 hours) obtained from the NCBI Gene Expression Omnibus (GEO), RNAseq and rMATS datasets of human RBP knockdowns as previously described [16, 38].

### AS events distribution patterns

The RNAseq datasets of the C57BL/6 mouse forebrain cover several developmental stages, namely E12.5, E13.5, E14.5, E15.5, E16.5, and P0. For each of these stages, two biological replicates were available. Using SUPPA2, AS events were initially pinpointed through pairwise comparisons across these time points. Differential alternative splicing events were then delineated based on criteria of | ΔPSI | > 0.1 and a p-value < 0.05.

Just as the PCA on gene expression data provides insights (as seen in Figure 2A), a PCA plot founded on PSI values also paints a coherent separation pattern for RNAseq samples across the various time points (illustrated in Figure 2B). A heatmap, constructed from these PSI values, effectively showcases notable shifts in AS events throughout the different developmental stages (Figure 2C). Concurrently, a volcano plot anchored in ΔPSI values offers a swift means to pinpoint statistically significant AS events (Figure 2D). The UpSet plot, on the other hand, serves to elucidate the intricate interplay between AS event sets. For instance, it highlights overlaps in differential AS events across the developmental timeline (as visualized in Figure 2E).

**Figure 2.**
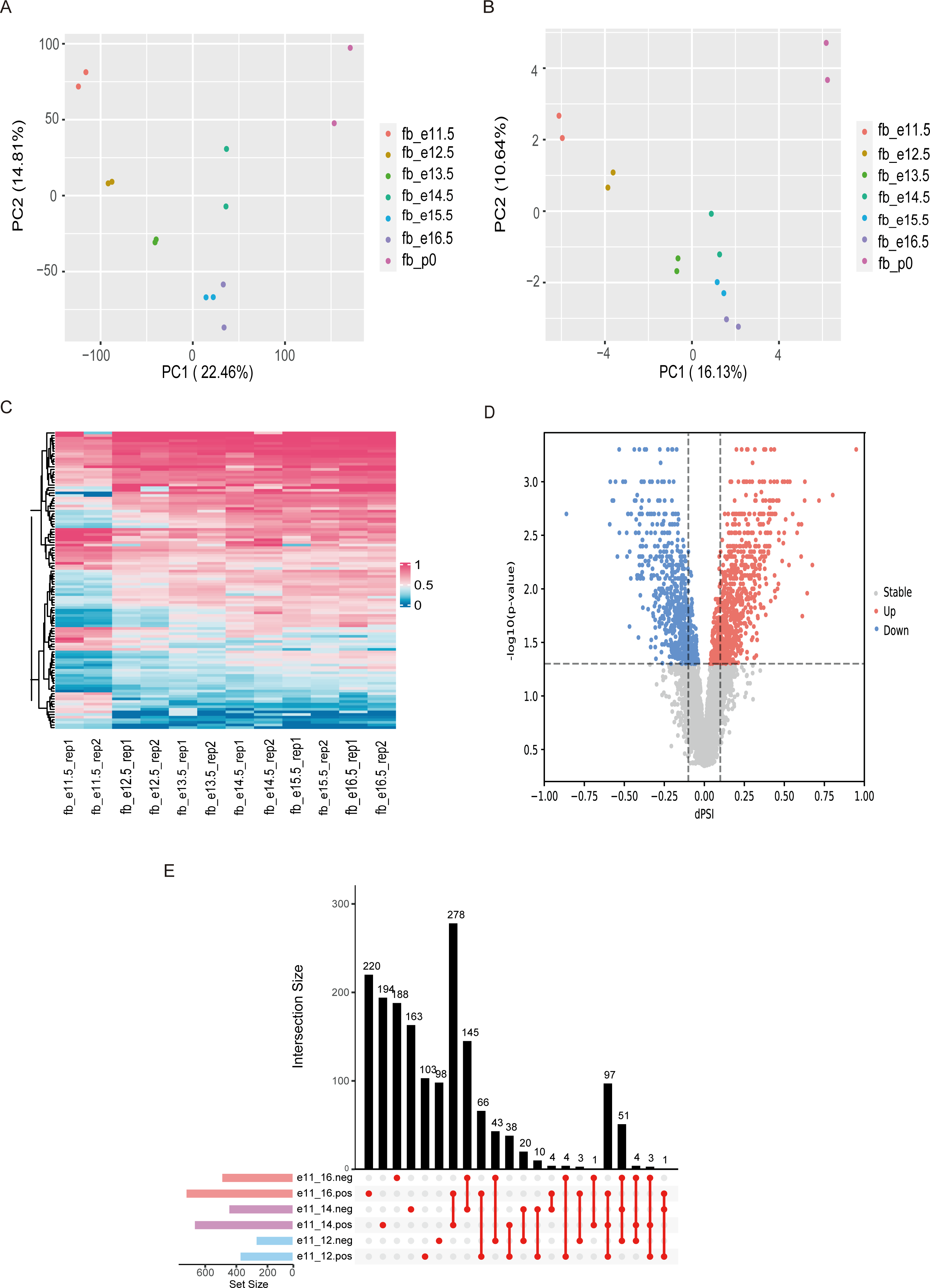
AS events distribution during embryonic mouse forebrain development: E11.5 - P0. **A.** PCA analysis based on gene expression data **B.** PCA analysis based on PSI values **C.** Heatmap on PSI values **D.** Volcano plot on ΔPSI values, neg: negative ΔPSI values, pos: positive ΔPSI values **E.** Upset plot for visualizing the intersection of different AS event sets

### Functional enrichment analysis

For our study, we selected differential AS genes during C57BL/6 mouse forebrain development (spanning E11.5 to P0) as test cases for gene-level functional enrichment analysis. A recurrent challenge with current enrichment analyses is the generation of redundant biological terms, complicating interpretation and potentially obscuring crucial biological events. To address this, we computed semantic similarities among enriched GO terms, clustering closely related ones. As a case in point, 538 significant GO terms were grouped into major clusters for the 761 differential AS genes identified between E11.5 and E16.5 (as shown in Figure 3A).

**Figure 3.**
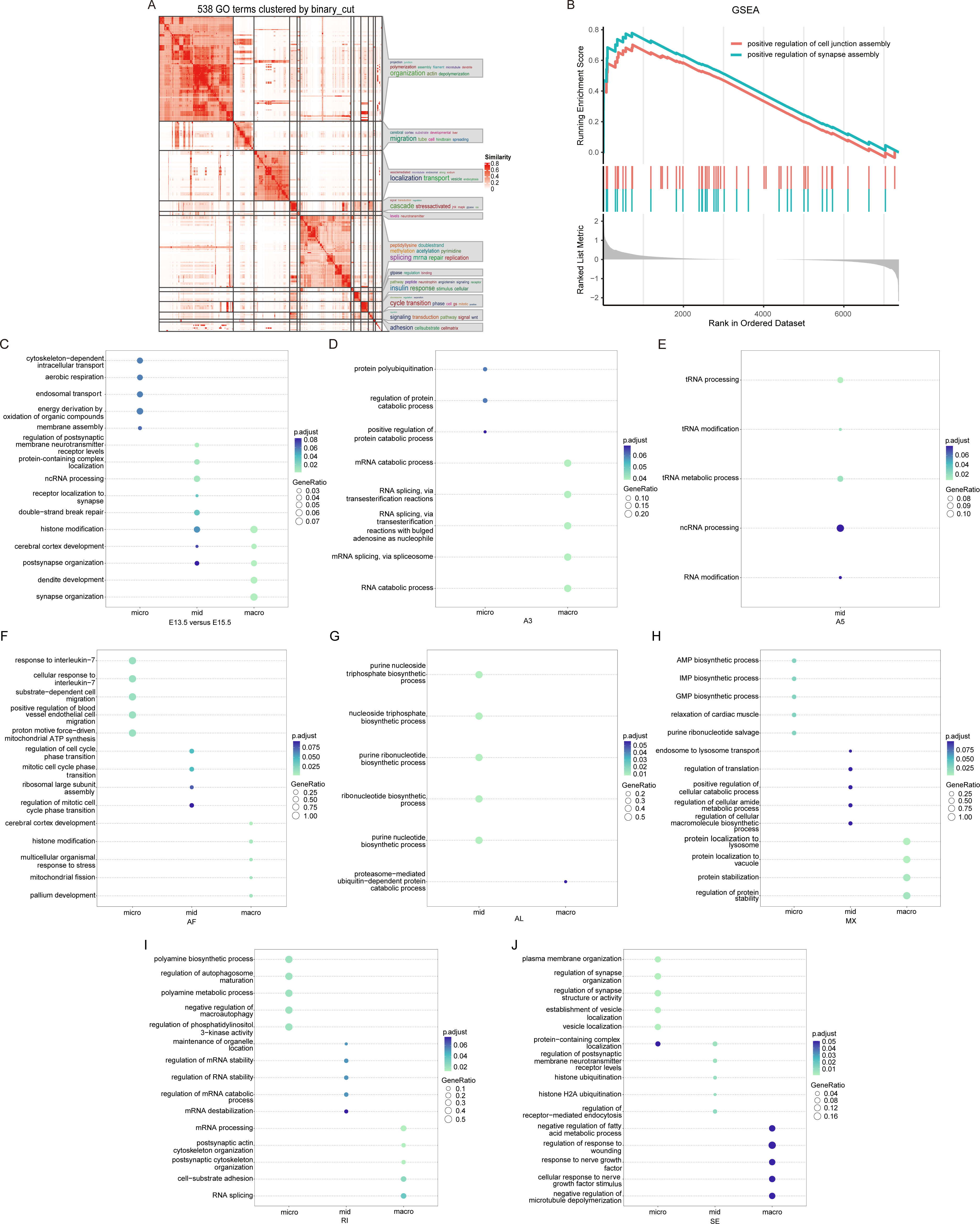
Functional enrichment analysis of differential AS events at gene-level before and after clustering. **A.** Clustering similar GO terms using simplifyEnrichment R package for differential AS genes identified between embryonic mouse forebrain E11.5 and E16.5 **B.** Gene-level GSEA analysis shows enriched pathways for AS genes with large ΔPSI values between embryonic mouse forebrain E11.5 and E16.5 **C.** Enriched GO terms for three groups of differential AS genes identified between embryonic mouse forebrain E13.5 and E15.5, which are clustered on the size of spliced exons **D-J.** GO enrichment analysis for clustered differential AS genes identified between embryonic mouse forebrain E13.5 and E15.5 after classification into seven types of AS: A3 (D), A5 (E), AF (F), AL (G), MX (H), RI (I), SE (J) GeneRatio: the number of enriched genes over the total number of input genes

However, solely relying on GO terms can exaggerate the significance of certain differentially expressed genes chosen based on arbitrary cutoffs. Utilizing both ORA and GSEA methods can provide a more comprehensive view. In Figure 3B, the GSEA result revealed a significant enrichment of AS genes with large ΔPSI values between E11.5 and E16.5 in processes like positive regulation of cell junction assembly and synapse assembly.

Prior to the functional enrichment analysis, we categorized differential AS genes into three groups based on spliced exon size. A recurring observation was the strong association of these groups with distinct functions. For instance, Figure 3C presents the GO enrichment results for differential AS genes between E13.5 and E15.5, categorized by exon size. When these genes are further classified into seven types of AS events and then clustered, the enriched GO pathways diverge distinctly (as visualized from Figure 3D-J). Even though at times differential AS genes have the same or similar enriched GO terms before classification, they appear to be functionally different after classification (Supplemental Figure 1). This trend is consistent, as demonstrated with differential AS genes identified between human stem cells 0h and 12h, which exhibit distinct GO terms before- and post-classification (Figures 4A-H).

**Figure 4.**
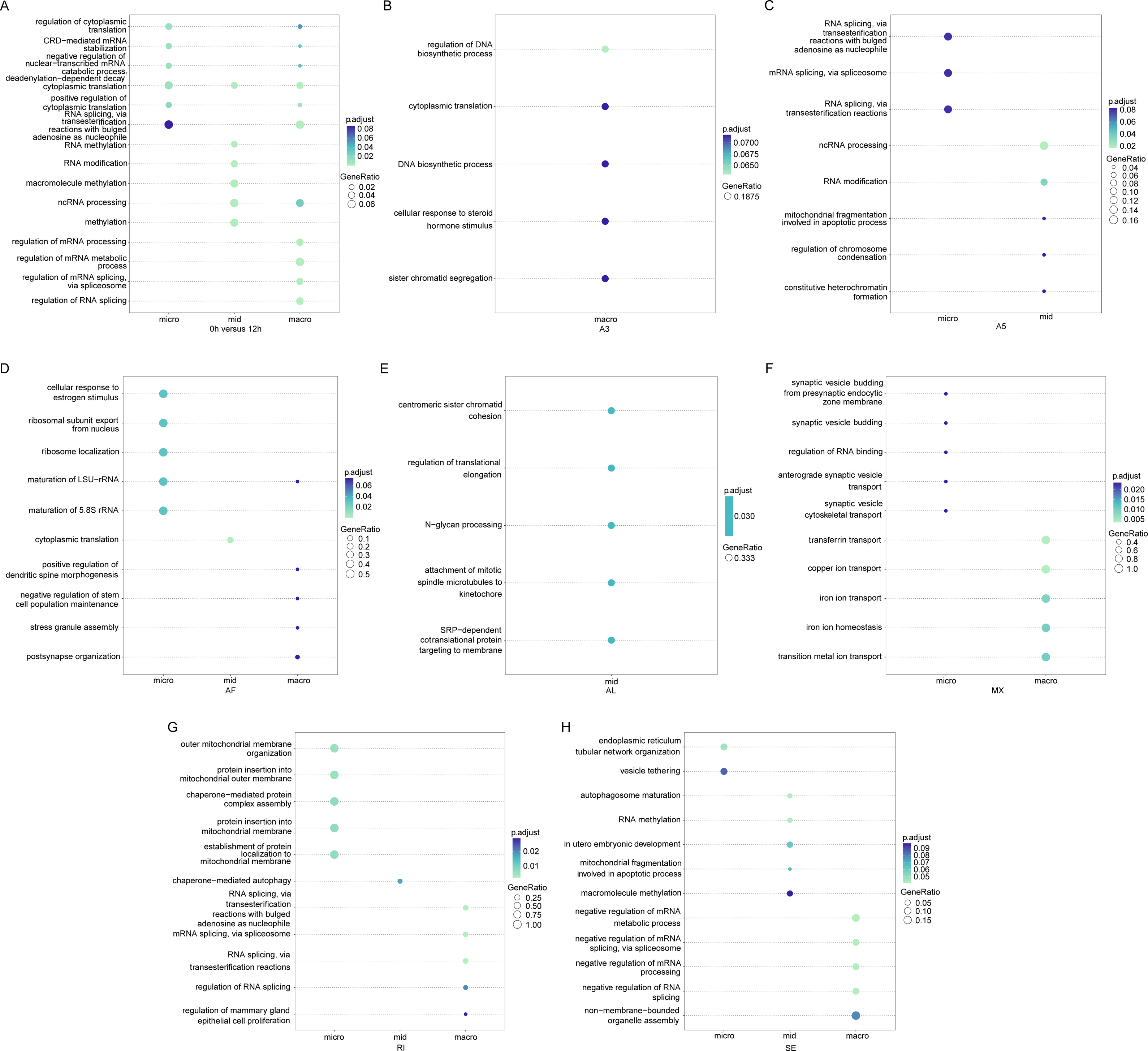
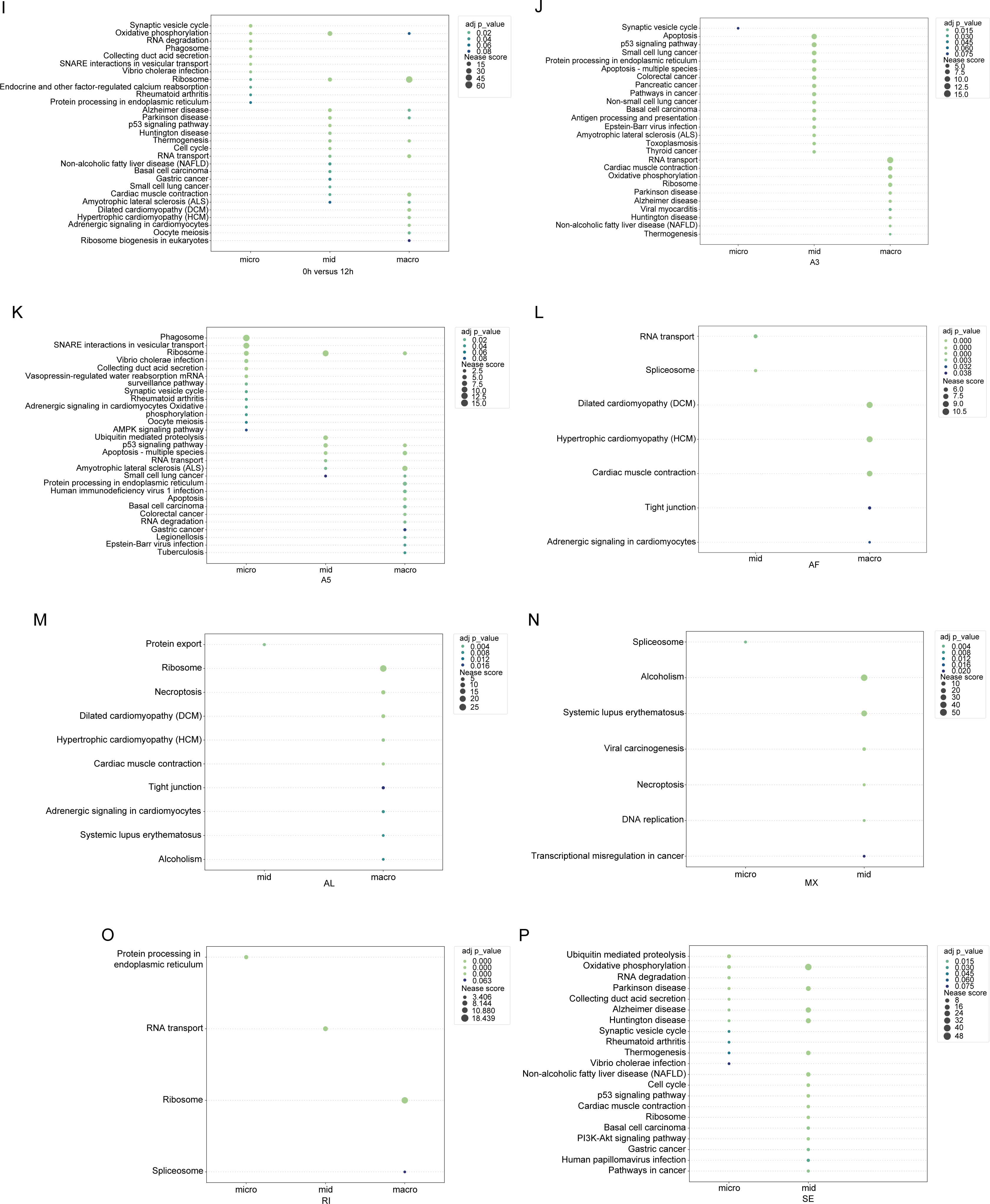
Functional enrichment analysis of differential AS events at gene- and exon-level before and after clustering. **A.** Enriched GO terms for three groups of differential AS genes identified between human stem cells 0h and 12h, which are clustered on the size of spliced exons **B-H.** GO enrichment analysis for clustered differential AS genes identified between human stem cells 0h and 12h after classification into seven types of AS: A3 (B), A5 (C), AF (D), AL (E), MX (F), RI (G), SE (H) **I.** Exon-level KEGG enrichment for three groups of spliced exons identified between human stem cells 0h and 12h **J-P.** Exon-level KEGG enrichment for three groups of spliced exons identified between human stem cells 0h and 12h after classification into types of AS: A3 (J), A5 (K), AF (L), AL (M), MX (N), RI (O), SE (P) GeneRatio: the number of enriched genes over the total number of input genes

However, it’s important to note that not every AS event has functional significance. Therefore, discerning meaningful structural and interactional changes in proteins caused by spliced exons can yield valuable biological insights into individual AS events or isoform shifts. Utilizing RNAseq datasets from human embryonic stem cells across seven neural differentiation time points (0, 3, 6, 12, 24, 48, and 72 h) [45], we spotlighted the functional consequences of spliced exons, complementing the gene-level enrichment analysis. For instance, Figure 4I-P presents the results of a KEGG enrichment analysis at the exon-level for differential AS genes identified between human stem cells 0h and 12h, showcasing distinct results for micro-, mid-sized, and macro-exons both pre- and post-classification.

While the above analyses were built on SUPPA2, we expanded our analysis to a larger dataset, the RNAseq of 185 RBP knockdowns, with AS events identified via rMATS. The results echoed our earlier findings. Supplemental Figure 2 illustrates the functional enrichment analysis of clustered AS genes and spliced exons in the SE type for six RBP knockdowns.

### RBP motif enrichment analysis and splicing effects

RBPs broadly regulate RNA transcripts at both transcriptional and post-transcriptional levels, such as splicing, modification, and translation [39, 40]. Utilizing data from a large-scale binding and functional study of human RBPs, we demonstrate that ASTK is adept at discerning the relationship between specific RBPs and splicing regulation. While a recently introduced tool, SpliceTools, was used to probe the influence of RBP knockdowns on the RI and SE types of AS events [16], ASTK widens the analysis scope to embrace all AS types.

Figures 5A-E provide a comprehensive panorama of the splicing effects triggered by individual RBP knockdowns. Beyond the commonly recognized splicing factors such as U2AF1 and U2AF2, our analysis unearthed potential key regulators for each AS type. To cite a few examples, PUF60 stands out for A3, SRSF1 and MAGOH shine in A5, SNRNP200 is notable for MX, while SF3B4 and TAF15 dominate in RI. For SE, SRSF1 and PUF60 emerged as prominent. Each of these knockdowns led to pronounced changes in AS events. Delving deeper, ASTK allowed us to elucidate the distinct regulatory roles of each RBP concerning exon inclusion or exclusion across various AS types. For instance, PUF60 knockdown predominantly triggers exon exclusion in SE (as seen in Figure 5F). In contrast, while SF3B4 knockdown encourages exon skipping in SE, it appears to promote intron inclusion in RI (Figure 5G). The underlying mechanisms for such outcomes remain a topic of future exploration. Our insights will undoubtedly pave the way for more in-depth functional characterization of these splicing factors.

**Figure 5.**
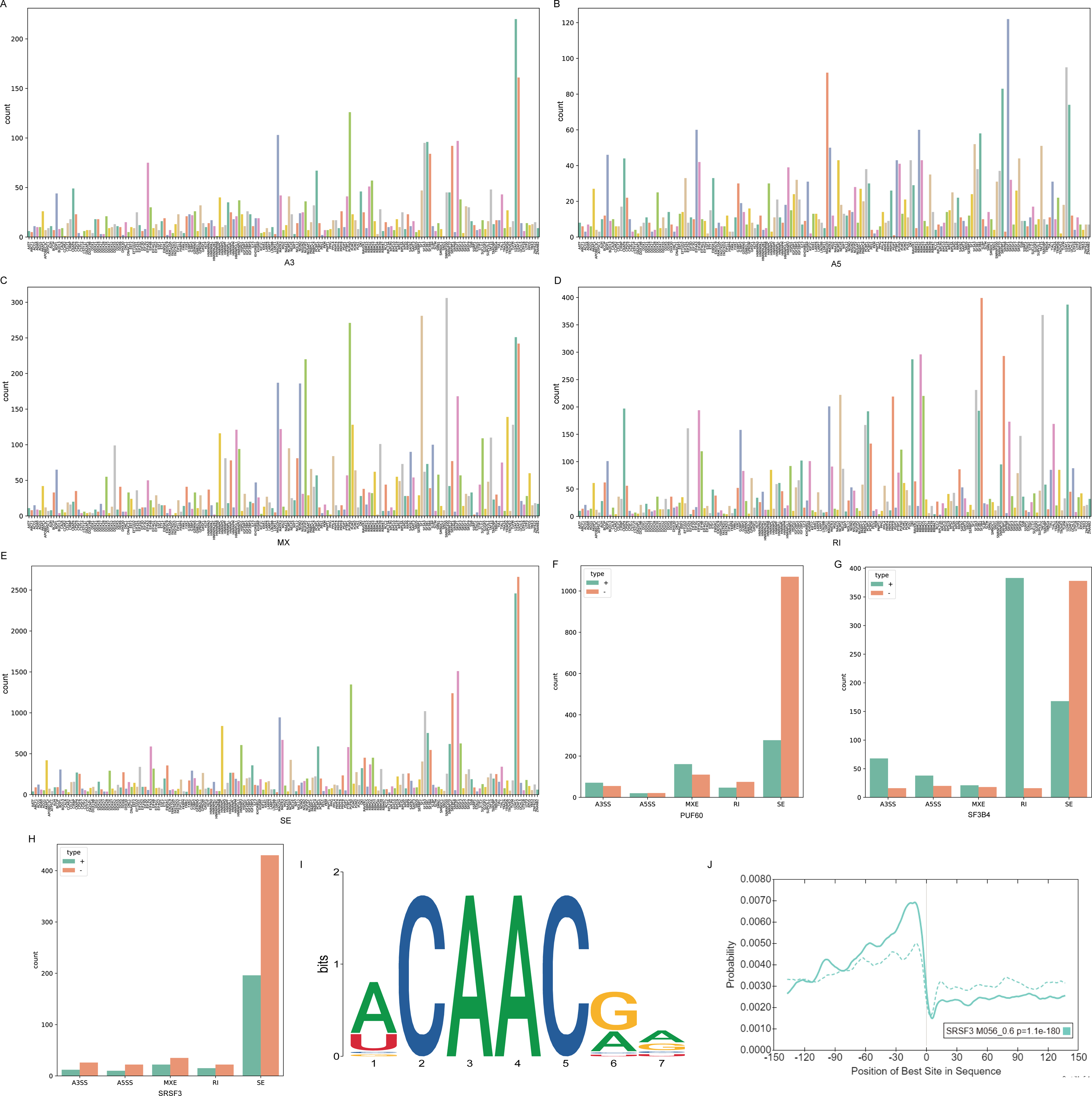
Splicing regulation by RBPs. **A.** Splicing changes in different types of AS caused by individual RBP knockdowns: A3 (A), A5 (B), MX (C), RI (D), SE (E). Y-axis (counts) - number of AS events; X-axis - each RBP knockdown **F.** PUF60 knockdown induces exon skipping. (+: exon inclusion; -: exon exclusion) **G.** SF3B4 knockdown promotes exon skipping and intron retention. (+: exon inclusion; -: exon exclusion) **H.** SRSF3 knockdown promotes exon skipping. (+: exon inclusion; -: exon exclusion) **I.** The sequence logo of the enriched SRSF3 binding motif **J.** The density curve of the SRSF3 binding motif in the flanking region of alternative 5’ splice site in SE for human embryonic stem cells at 72 hours, solid line – high PSI group, dashed line – low PSI group

The manner in which RBPs regulate AS is intrinsically linked to their binding site preferences and positions adjacent to the splice sites. Taking human embryonic stem cells at the 72-hour mark as a case in point, Figure 5I illustrates the sequence logo for the RBP, SRSF3, derived from the CISBP-RNA motif database [41]. It’s evident that SRSF3’s binding affinity in the flanking exon of the alternative 5’ splice site is markedly elevated in high PSI groups compared to their low PSI counterparts in SE (Figure 5J). Prior research has highlighted that SRSF3 binding sites predominantly enriched in the exonic region close to the splice sites [42], which is essential for exon inclusion. SRSF3 knockdown has been shown to stimulate the selection of alternative 5’ splice sites and foster exon skipping in human cell lines [42]. Our analysis of RBP knockdowns corroborates these observations, especially in the context of SE (Figure 5H).

In summation, our results underscore the potential of ASTK as a valuable tool for researchers, enabling them to glean critical insights into the regulatory roles of RBPs or splicing factors in the realm of splicing regulation.

### Specific sequence features and epigenetic marks associated with AS

In eukaryotic cells, the majority of AS events occur co-transcriptionally, which are highly sensitive to sequence features, RNA polymerase II elongation rate, and chromatin modifications and structure [1, 28, 34, 36, 43, 44]. Consequently, constructing an optimal splicing code demands the integration of these diverse features, operating synergistically across multiple levels.

Our examination utilized the ENCODE datasets, spanning RNAseq, ChIPseq, and ATACseq during mouse forebrain development from E12.5 to E16.5. This exploration encompassed eight critical histone modifications and seven types of AS events, computed with SUPPA2 (Figure 1B). The features, spanning sequence elements, ATAC signals, and ChIPseq signals of histone modifications, were extracted systematically, covering all AS classes over five developmental stages.

The sequence context around the splice sites from both constitutive and alternative exons is derived to examine potential patterns associated with AS. Upstream and downstream sequences from each exon junction are extracted: +/- 150 nt flanking region surrounding 5’ splice site, +/- 150 nt flanking region surrounding 3’ splice site. After removing the conserved 5’ and 3’ splice site sequences, the GC content in each flanking region is calculated. Lengths of related exons and introns for each AS event are computed.

Sequence features are part of genomic architecture, contributing to splice site and splicing unit recognition and affecting the splicing process [45, 46]. The GC content, integral to genomic organization, was found to influence the stability of DNA and mRNAs. We found that flanking regions surround alternative splice sites in AF and SE classes exhibited higher GC content in high PSI groups compared to their low PSI counterparts, suggesting a higher likelihood of selection for splice sites with elevated GC content (Figure 6A-B).

**Figure 6.**
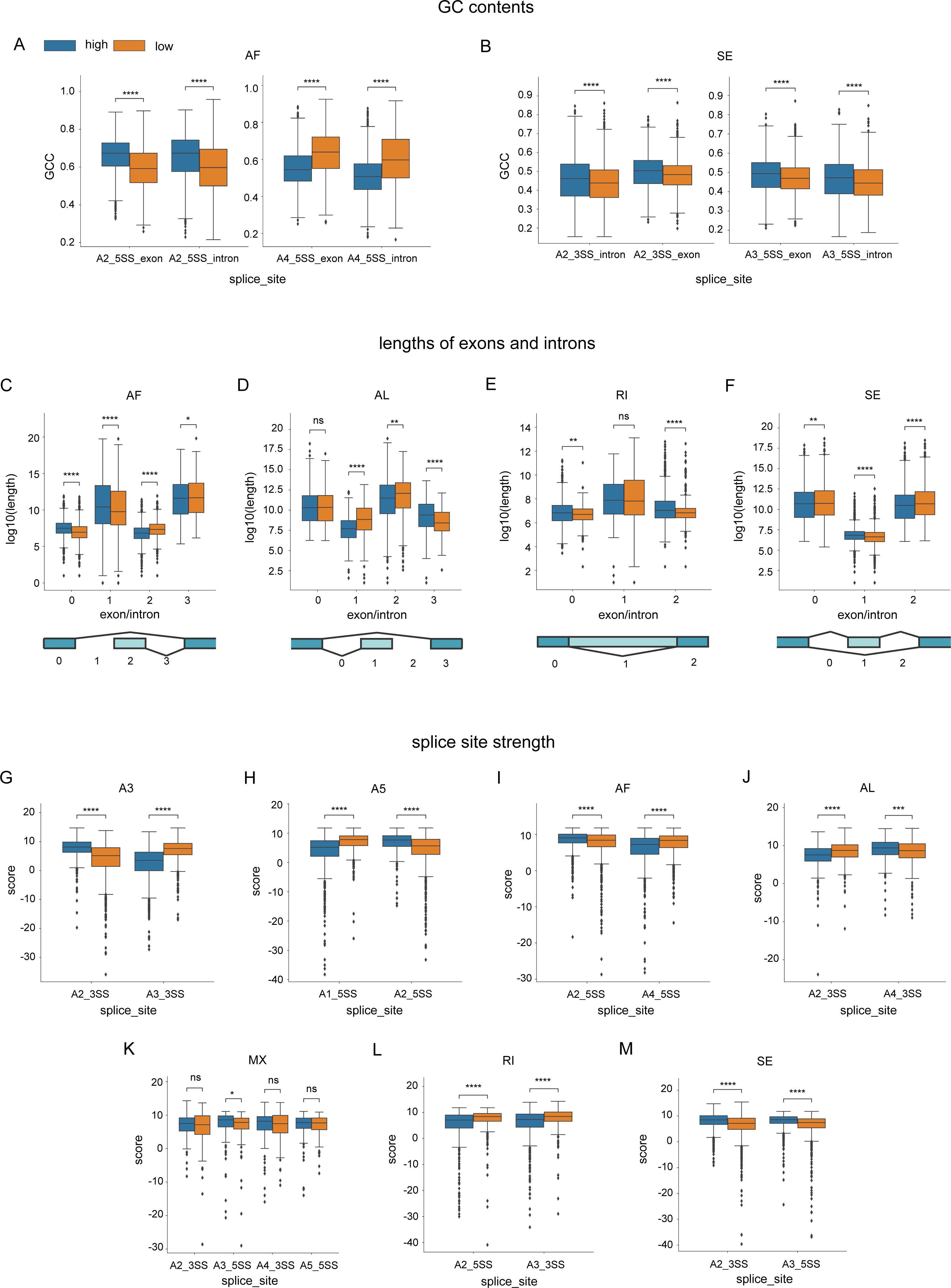
Sequence features associated with AS events at E16.5. **A-B.** +/- 150 nt flanking regions of each 5’ or 3’ splice site were extracted, and their GC contents were computed after removing the conserved 5’ or 3’ splice site sequences. In AF (A) and SE (B) classes, the high PSI group have a higher GC content in these flanking regions than the low PSI group. In addition, “_exon” and “_intron” suffix denote the exonic and intronic flanking regions of the splice site, respectively. **C-F.** Lengths of up- or downstream exons/ introns for each alternative splice site, converted to log base 2 values. In AF, AL, and ES classes (C, D, F), larger alternative exons are more likely to be spliced in than smaller ones. In RI (E), the PSI high group has significantly larger upstream and downstream exons of a retained intron than the PSI low group. **G-M.** Splice site strength is computed for each alternative splice site based on MaxEntScan. Higher splice site strength favors the inclusion of alternative exons in A3, A5, AF, AL, and ES (G-J, M). In contrast, lower spice site strength promotes the intron retention in RI (L).

We further delved into the influence of intron and exon lengths on splicing. Our observations underscored that in AF, AL, and SE events, larger alternative exons are more frequently spliced in (Figure 6C, D, F). Conversely, the lengths of retained introns in the RI showed significant variation across PSI groups. Still, the flanking exons of a retained intron were markedly larger in the PSI high group (Figure 6E).

The selection of alternative splice sites is inherently competitive and depends on a number of factors. Splice site strength is one of them. We observed that the splice site strengths are significantly different between the PSI high versus its corresponding low groups for all AS classes except MX. In general, we found that the splice site strength is positively correlated with exon inclusion levels of 5 types of AS events except MX and RI (Figure 6G-J, M). For instance, for SE events, 5’ and 3’ splice sites flanking the skipped exons in the high PSI group appear to have significantly higher splice site strength than its corresponding low PSI group. Since PSI refers to the percent spliced-in value, our results suggest that given two 5’ or 3’ splice sites, the higher-scoring ones have a higher probability of being selected (Figure 6M). Interestingly, for RI, the PSI high group has lower strength in both 5’ and 3’ splice sites flanking the retained intron than its corresponding PSI low group (Figure 6L), indicating that these retained introns probably have weak splice sites flanked so that RNA polymerase II can read through and include the intron in the mRNA.

Our assessment also encompassed the integration of epigenetic features with AS events. For the ATAC signal and each histone modification, we summarized the ChIPseq signal in the ± 150 nt vicinity of each alternative splice site and its corresponding constitutive splice site. ATACseq is a popular method used to measure genome-wide chromatin accessibility. ATAC signal abundance is highly indicative of active transcription. We found that an increased ATAC signal is significantly associated with the alternative first exon selection in AF events (Figure 7A). Moreover, histone modifications, known regulators of gene expression [47]. have been posited to influence splicing outcomes [32], likely through a combinatorial manner [48]. Our study scrutinized the signal distributions of eight histone marks in regions adjacent to splice sites over five developmental stages. The patterns revealed varied enrichment of histone marks across AS classes and time points. For instance, in AF events, specific histone marks, such as H3K4me2, H3K4me3, H3K9ac, and H3K27ac, were observed to favor the inclusion of an alternative first exon, whereas others like H3K4me1, H3K27me3, and H3K36me3 demonstrated the opposite effect (Figure 7B-I). Meanwhile, no clear association was observed for H3K9me3 (Figure 7F). In SE events, H3K36me3 positively correlates with the PSI high groups (Figure 7R). However, H3K4me1, H3K4me2, and H3K27me3 appear to have negative correlations (Figure 7K-L, Q).

**Figure 7.**
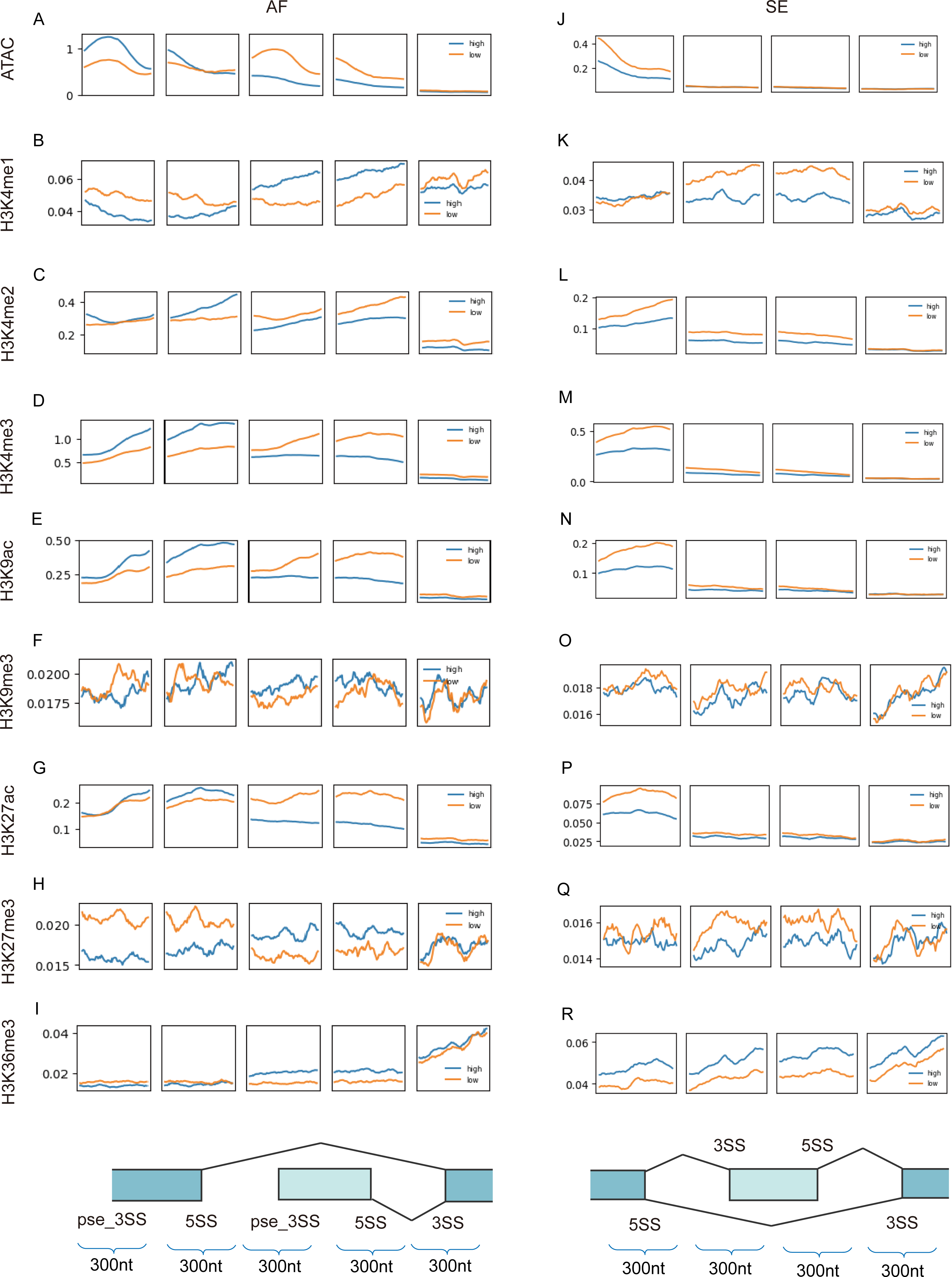
Epigenetic features associated with AS events at E16.5. **A-G.** In AF, several epigenetic features are positively correlated with the PSI high group: ATAC signal, H3K4me2, H3K4me3, H3K9ac, and H3K27ac. Splice sites are illustrated at the bottom. **H-N.** In SE, H3K36me3 has a positive correlation with the PSI high groups, while H3K4me1 and H3K27me3 have negative correlations. Splice sites are illustrated at the bottom.

### Model predictions and feature importance using AS events identified by SUPPA2

To uncover how sequence and epigenetic features influence distinct types of AS events, we utilized a random forest binary classifier. This classifier differentiated each AS type into PSI high versus PSI low groups, using both sequence and epigenetic features. Model performance was gauged using accuracy and AUC (area under the ROC curve) metrics, based on a 5-fold cross-validation process (Figure 8).

**Figure 8.**
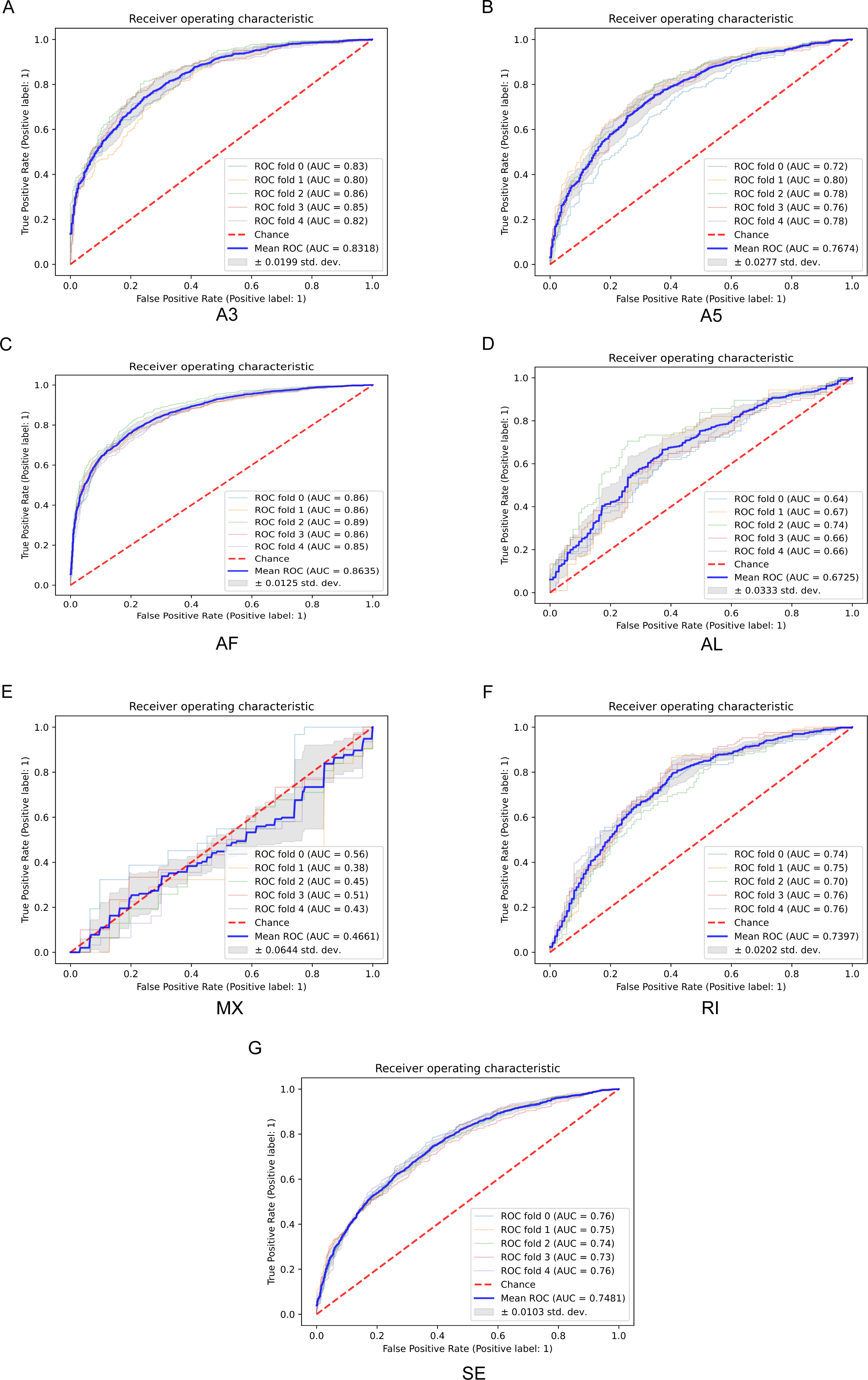
ROC curves of PSI high versus low classifications for seven AS types at E16.5 using pooled sequence and epigenetic features. **A-G.** ROC was measured with 5-fold cross-validation for each AS type. The mean ROC AUC scores are reported: 0.83 (A3), 0.77 (A5), 0.86 (AF), 0.67 (AL), 0.47 (MX), 0.74 (RI), and 0.75 (SE).

Remarkably, sequence features alone - which encompass the GC content around splice sites, lengths of spliced exons, adjacent introns, and splice site strength - displayed commendable predictive capability for all AS classes, with MX being an exception. As shown in Table 1, when predicting PSI high (top 25%) versus low (bottom 25%) using combined sequence features, the model accuracy varied significantly for seven AS classes across different developmental stages: A3 (0.75-0.77), AF (0.74-0.76), A5 (0.71-0.72), RI (0.69-0.70), SE (0.68-0.69), AL (0.66-0.70), MX (0.44-0.56). AUC scores are in line with accuracy values, A3 (0.84-0.85), AF (0.82-0.83), A5 (0.78-0.79), RI (0.76-0.77), SE (0.74-0.76), AL (0.70-0.74), MX (0.47-0.59).

**Table 1.** Accuracies (ACC) and AUC values of random forest models to predict PSI high (top 25%) versus low groups (bottom 25%) for seven AS types at each developmental time point using combined sequence features as input (**AS data generated by SUPPA2**)

|  | e12.5 |  | e13.5 |  | e14.5 |  | e15.5 |  | e16.5 |  |
| --- | --- | --- | --- | --- | --- | --- | --- | --- | --- | --- |
|  | ACC | AUC | ACC | AUC | ACC | AUC | ACC | AUC | ACC | AUC |
| A3 | 0.7597 | 0.8430 | 0.7597 | 0.8453 | 0.7695 | 0.8532 | 0.7546 | 0.8421 | 0.7532 | 0.8410 |
| A5 | 0.7077 | 0.7794 | 0.7165 | 0.7869 | 0.7232 | 0.7911 | 0.7159 | 0.7909 | 0.7168 | 0.7825 |
| AF | 0.7585 | 0.8333 | 0.7424 | 0.8170 | 0.7464 | 0.8254 | 0.7466 | 0.8249 | 0.7469 | 0.8246 |
| AL | 0.6990 | 0.7402 | 0.6606 | 0.6987 | 0.6800 | 0.7284 | 0.6692 | 0.7183 | 0.6606 | 0.7110 |
| MX | 0.5610 | 0.5928 | 0.5286 | 0.5573 | 0.5068 | 0.5014 | 0.4443 | 0.4731 | 0.5324 | 0.5372 |
| RI | 0.6908 | 0.7569 | 0.6951 | 0.7629 | 0.6962 | 0.7554 | 0.6963 | 0.7642 | 0.7001 | 0.7651 |
| SE | 0.6912 | 0.7586 | 0.6834 | 0.7467 | 0.6753 | 0.7431 | 0.6798 | 0.7466 | 0.6768 | 0.7414 |

Subsequently, we investigated the effects of eight histone modifications and chromatin accessibility on AS. We found that epigenetic factors significantly impact how the first alternative exon is selected and how the alternative exon is skipped (Table 3). The prediction accuracy on the PSI high versus its corresponding PSI low group using these epigenetic features is approximately 0.77 and 0.59 for AF and SE, respectively. Taking one step further, we ran the model with pooled sequence and epigenetic features. Combining both sequence and epigenetic features only slightly enhanced the predictive accuracy for AF (Table 5).

To reveal the regulatory mechanisms of AS, we were in search of features with major contributions to the classification. In our random forest model, variable importance is quantified by Gini importance. Consequently, using E16.5 as an example, we obtained a ranked feature importance list for every AS class (Figure 9). One of the major findings is that seven AS classes exhibit distinctive regulatory features. Since the accuracy of MX is fluctuating around 50%, a random guessing level, we focused on the other six types of AS events. The results indicate that splice site strength is the single most important factor to determine the PSI high versus low for A3 and A5 classes. While, multiple sequence and epigenetic features significantly contribute to the classification in AF, including GC contents, the lengths of introns and exons, splice site strength, ATAC signal, H3K4me3, H3K9ac, H3K27ac, and H3K4me2. Like AF, sequence features and specific epigenetic features are strongly associated with SE classification. In AL, only two top features were identified: the lengths of introns and exons, and splice site strength. Moreover, sequence features significantly influence the classification in RI (Table 7).

**Figure 9.**
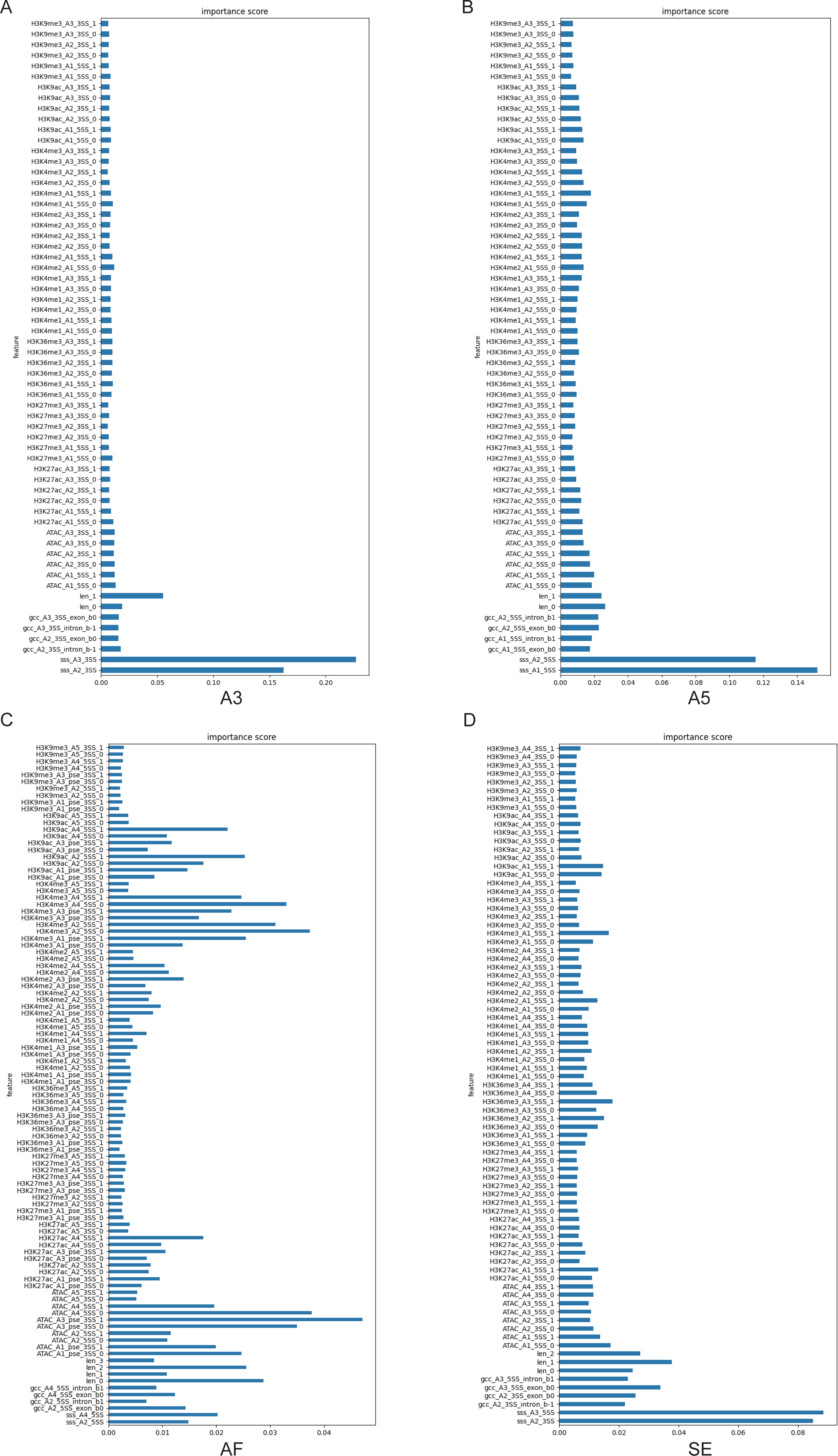
Feature importance in A3, A5, AF, and SE. **A-B.** In A3 and A5, the splice site strength is the most critical feature deciding PSI high versus low classification. **C-D.** In AF and SE, multiple sequence features and epigenetic factors strongly influence PSI high versus low classification.

We further assessed the model’s sensitivity to chosen cutoff values, such as the top 33% and bottom 33% (data not shown), as well as the top 40% and bottom 40%. The results remained largely consistent across varying cutoffs. However, accuracy experienced a minor dip with increased cutoffs due to reduced sensitivity (Tables 2, 4, 6, and 8).

**Table 2.** Accuracies (ACC) and AUC values of random forest models to predict PSI high (top 40%) versus low groups (bottom 40%) for seven AS types at each developmental time point using combined sequence features as input (**AS data generated by SUPPA2**)

|  | e12.5 |  | e13.5 |  | e14.5 |  | e15.5 |  | e16.5 |  |
| --- | --- | --- | --- | --- | --- | --- | --- | --- | --- | --- |
|  | ACC | AUC | ACC | AUC | ACC | AUC | ACC | AUC | ACC | AUC |
| A3 | 0.7208 | 0.7934 | 0.7178 | 0.7926 | 0.7185 | 0.7983 | 0.7156 | 0.7905 | 0.7113 | 0.7920 |
| A5 | 0.6744 | 0.7309 | 0.6622 | 0.7232 | 0.6670 | 0.7273 | 0.6652 | 0.7300 | 0.6612 | 0.7274 |
| AF | 0.7009 | 0.7681 | 0.6881 | 0.7545 | 0.7029 | 0.7692 | 0.6940 | 0.7617 | 0.6934 | 0.7621 |
| AL | 0.6456 | 0.6906 | 0.6273 | 0.6694 | 0.6526 | 0.6962 | 0.6478 | 0.6785 | 0.6326 | 0.6811 |
| MX | 0.5191 | 0.5174 | 0.5021 | 0.5008 | 0.4935 | 0.5099 | 0.4629 | 0.4640 | 0.5282 | 0.5379 |
| RI | 0.6414 | 0.7055 | 0.6440 | 0.7083 | 0.6342 | 0.6883 | 0.6647 | 0.7119 | 0.6469 | 0.7062 |
| SE | 0.6663 | 0.7231 | 0.6570 | 0.7102 | 0.6551 | 0.7106 | 0.6552 | 0.7121 | 0.6540 | 0.7101 |

**Table 3.**
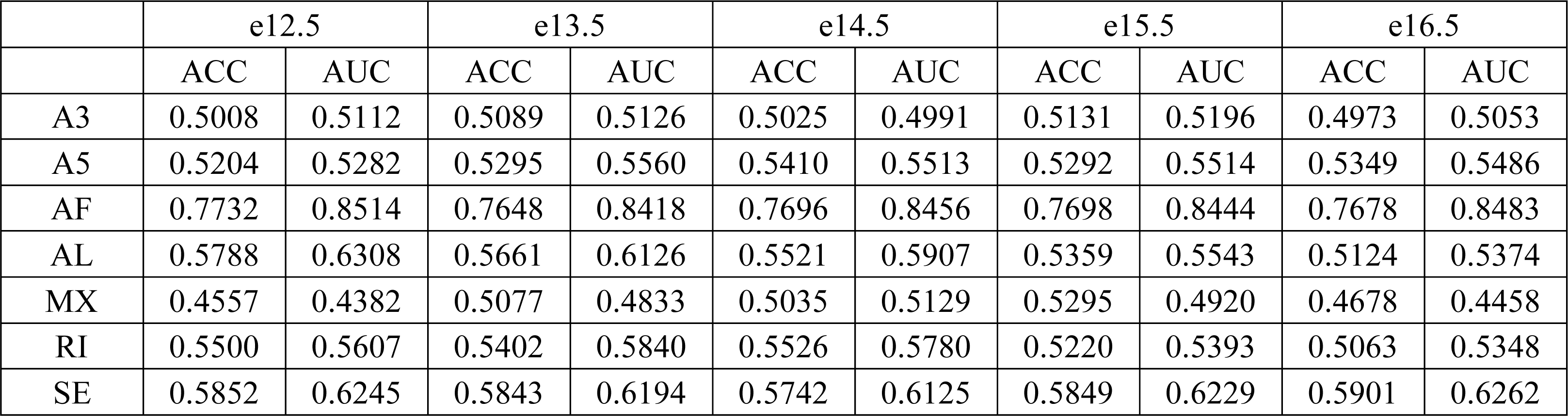
Accuracies (ACC) and AUC values of random forest models to predict PSI high (top 25%) versus low groups (bottom 25%) for seven AS types at each developmental time point using combined epigenetic features as input (**AS data generated by SUPPA2**)

**Table 4.**
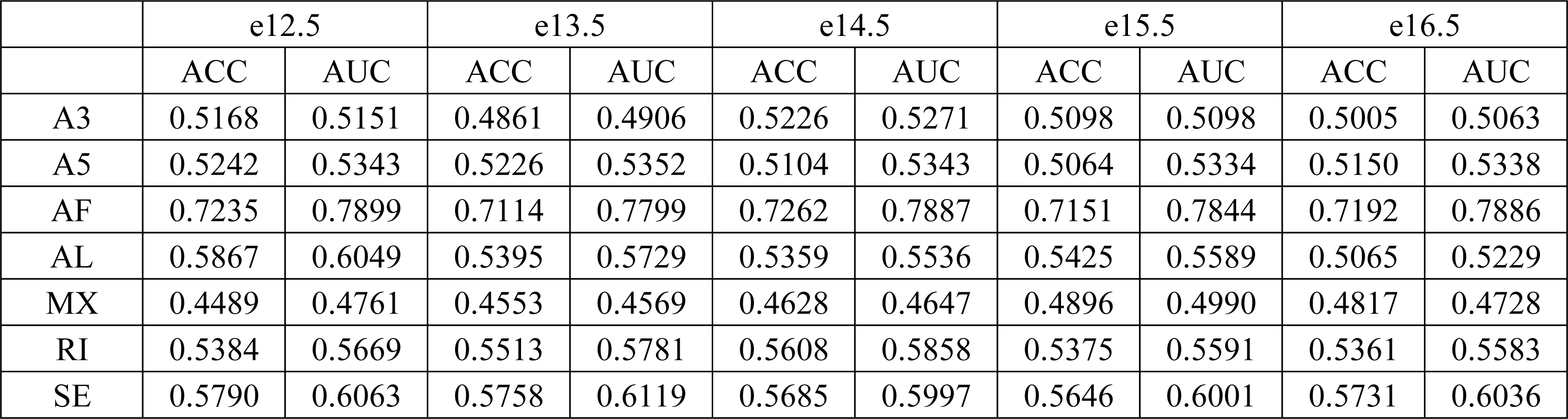
Accuracies (ACC) and AUC values of random forest models to predict PSI high (top 40%) versus low groups (bottom 40%) for seven AS types at each developmental time point using combined epigenetic features as input (**AS data generated by SUPPA2**)

**Table 5.** Accuracies (ACC) and AUC values of random forest models to predict PSI high (top 25%) versus low groups (bottom 25%) for seven AS types at each developmental time point using pooled sequence and epigenetic features as input (**AS data generated by SUPPA2**)

|  | e12.5 |  | e13.5 |  | e14.5 |  | e15.5 |  | e16.5 |  |
| --- | --- | --- | --- | --- | --- | --- | --- | --- | --- | --- |
|  | ACC | AUC | ACC | AUC | ACC | AUC | ACC | AUC | ACC | AUC |
| A3 | 0.7441 | 0.8286 | 0.7537 | 0.8336 | 0.7629 | 0.8420 | 0.7473 | 0.8266 | 0.7437 | 0.8318 |
| A5 | 0.7027 | 0.7693 | 0.6949 | 0.7714 | 0.7052 | 0.7794 | 0.7113 | 0.7713 | 0.7013 | 0.7674 |
| AF | 0.7918 | 0.8716 | 0.7892 | 0.8599 | 0.7828 | 0.8628 | 0.7823 | 0.8622 | 0.7771 | 0.8635 |
| AL | 0.6625 | 0.7353 | 0.6342 | 0.6791 | 0.6411 | 0.7017 | 0.6257 | 0.6765 | 0.6398 | 0.6725 |
| MX | 0.5134 | 0.5052 | 0.5071 | 0.4956 | 0.5137 | 0.5065 | 0.4507 | 0.4272 | 0.4805 | 0.4661 |
| RI | 0.6900 | 0.7333 | 0.6692 | 0.7288 | 0.6822 | 0.7371 | 0.6691 | 0.7288 | 0.6756 | 0.7397 |
| SE | 0.6923 | 0.7602 | 0.6828 | 0.7475 | 0.6773 | 0.7393 | 0.6858 | 0.7490 | 0.6776 | 0.7481 |

**Table 6.** Accuracies (ACC) and AUC values of random forest models to predict PSI high (top 40%) versus low groups (bottom 40%) for seven AS types at each developmental time point using pooled sequence and epigenetic features as input (**AS data generated by SUPPA2**)

|  | e12.5 |  | e13.5 |  | e14.5 |  | e15.5 |  | e16.5 |  |
| --- | --- | --- | --- | --- | --- | --- | --- | --- | --- | --- |
|  | ACC | AUC | ACC | AUC | ACC | AUC | ACC | AUC | ACC | AUC |
| A3 | 0.7054 | 0.7796 | 0.7135 | 0.7821 | 0.7150 | 0.7865 | 0.7093 | 0.7823 | 0.7101 | 0.7843 |
| A5 | 0.6551 | 0.7149 | 0.6514 | 0.7074 | 0.6488 | 0.7087 | 0.6503 | 0.7105 | 0.6536 | 0.7172 |
| AF | 0.7348 | 0.8053 | 0.7281 | 0.7967 | 0.7286 | 0.8018 | 0.7274 | 0.8000 | 0.7291 | 0.8029 |
| AL | 0.6186 | 0.6737 | 0.5910 | 0.6421 | 0.6208 | 0.6791 | 0.6229 | 0.6579 | 0.6207 | 0.6758 |
| MX | 0.4732 | 0.4785 | 0.4686 | 0.4738 | 0.5308 | 0.5195 | 0.4731 | 0.4516 | 0.5201 | 0.5211 |
| RI | 0.6606 | 0.7008 | 0.6408 | 0.7018 | 0.6462 | 0.6840 | 0.6516 | 0.7029 | 0.6533 | 0.6946 |
| SE | 0.6669 | 0.7282 | 0.6534 | 0.7112 | 0.6586 | 0.7122 | 0.6561 | 0.7147 | 0.6602 | 0.7144 |

**Table 7.**
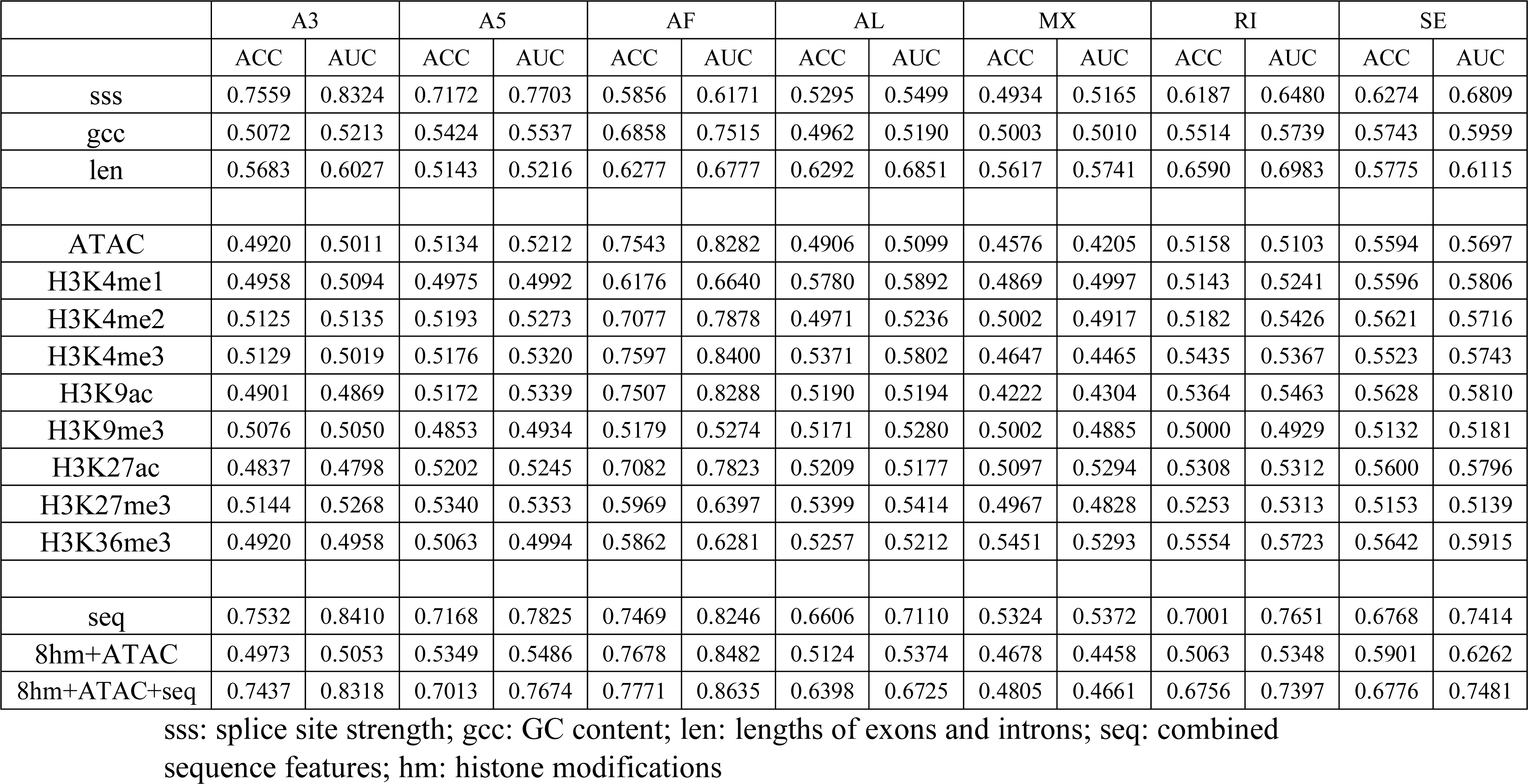
Accuracies (ACC) and AUC values of random forest models to predict PSI high (top 25%) versus low groups (bottom 25%) for seven AS types at E16.5 using each or combined sequence and epigenetic features as input (**AS data generated by SUPPA2**)

**Table 8.**
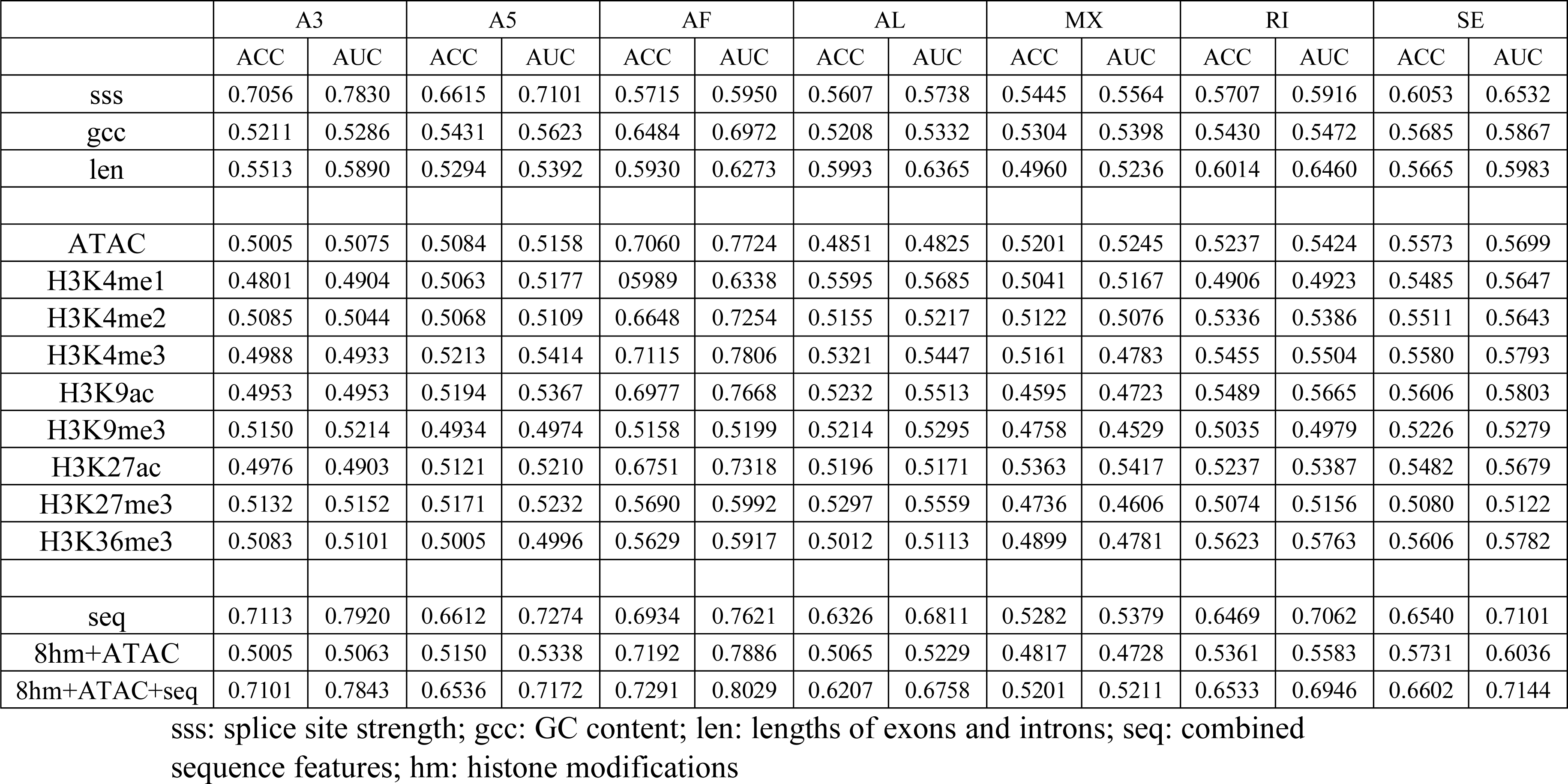
Accuracies (ACC) and AUC values of random forest models to predict PSI high (top 40%) versus low groups (bottom 40%) for seven AS types at E16.5 using each or combined sequence and epigenetic features as input (**AS data generated by SUPPA2**)

### Model predictions and feature importance using AS events identified by rMATS

Further validation of our machine learning models was undertaken using another benchmarking tool, rMATS. Recognized as a popular tool, rMATS identifies five types of splicing events: A3, A5, MX, RI, and SE. Performance assessment of the model was executed across five time points: E12.5, E13.5, E14.5, E15.5, and E16.5. For each AS type, the top 25% and bottom 25% of the PSI data generated by rMATS were delineated as PSI high and low, respectively. The prediction accuracies exhibited similarities to the outcomes derived from SUPPA2 output files. Specifically, the accuracies are as follows: A3 (0.74-0.75), A5 (0.75-0.77), RI (0.74-0.76), SE (0.67-0.68), MX (0.60-0.63) (Supplemental Table 1). ROC curves were employed to gauge model performance (Supplemental Figure 3).

Focusing on E16.5 as a representative time point, it was discerned that sequence features predominantly dictate the classifications for A3, A5, and RI. In contrast, SE’s classification is shaped by a combination of multiple sequence and epigenetic features (for details, see Supplemental Table 2 and Supplemental Figure 4).

In essence, the model’s validation using both benchmarking tools, rMATS and SUPPA2, yielded consistent results, underscoring the robustness of our approach.

## Discussion

The advancement of sequencing technologies has paved the way for expansive transcriptomic analyses. One of the paramount applications of RNAseq in recent years has been genome-wide profiling of alternative splicing, leading to the creation of comprehensive databases like MeDAS and ASCancer [49, 50]. As a core gene regulatory process, AS plays a pivotal role in sustaining cellular physiology and behavior. Conversely, aberrations in AS are linked with a myriad of human diseases and cancers [17]. Studies even suggest that nearly 20% of human disease-related mutations have an impact on splicing [51].

This underscores the importance of AS research, as it can significantly enhance our understanding of gene regulation, disease causation, and potentially aid in the creation of AS-targeted therapies. Our endeavor through this study was to craft an exhaustive pipeline to unravel the functional and regulatory facets of AS. While several computational tools have emerged to quantify transcript isoforms and identify splicing events [3, 4, 7, 8, 12], the functional elucidation of AS events remains an intricate challenge. Traditional gene-level enrichment analyses like GO, KEGG, and GSEA primarily target differential AS genes, often overlooking the fact that a single gene can harbor multiple AS events and exhibit diverse splicing patterns. Recognizing this gap, we incorporated the NEASE method in our software to specifically probe the functional repercussions at the exon level. Furthermore, previous work has found that spliced exons less than 250 nt are significantly conserved in terms of both length and sequence across vertebrate species [18, 25, 52, 53]. Our classification of differential AS genes and exons based on exon lengths led to intriguing discoveries regarding their distinct functional inclinations.

The intricate nature of AS reflects the multifaceted landscape of gene regulation. Genomic architecture, including splice site strength, GC content, and exon-intron structure, lays the foundation for the splicing unit. Concurrently, epigenetic elements, from histone modifications to chromatin structures, wield influence over exon splicing by modulating the RNA polymerase II elongation rate and aiding in the recruitment of splicing machinery [31, 32, 36, 44, 48, 54, 55]. The culmination of these processes is orchestrated by splicing factors that position themselves at specific binding sites to facilitate splicing [1, 29]. A recent study shows that AS genes shared with specific histone marks tend to have similar RBP binding motifs and enrich in common functional pathways [48], suggesting a strong functional coupling between epigenomic features and coordinately regulated cell-specific alternative splicing. Given the complex genomic structures and plethora of splicing regulators, decoding the chromatin signatures and regulatory features associated with AS becomes computationally arduous. To address this, our ASTK is equipped with modules to proficiently identify sequence features, RBP binding motifs, and other pertinent epigenomic signatures.

Crucially, our incorporation of machine learning algorithms into ASTK has enabled us to demystify the regulatory codes governing various AS classes. Our models, trained on sequence and epigenetic features, revealed distinctive predictive features for different AS classes. Splice site strength emerged as a predominant predictor for A3 and A5, while other features like ATAC signal, H3K4me3, and H3K9ac showcased significant predictive power for AF. This suggests that each AS class might be governed by unique regulatory mechanisms with varying contributions from sequence and epigenetic features.

To the best of our understanding, ASTK is the inaugural platform amalgamating machine learning for a comprehensive analysis of AS events. Through rigorous testing on experimental multi-omics datasets, we have demonstrated the efficacy and potential of ASTK in advancing AS research, offering valuable insights into its regulation, functional implications, and evolutionary trajectory.

## Acknowledgments

We thank Dr. Jianming Zeng (University of Macau), and all the members of his bioinformatics team, and biotrainee, for generously sharing their experience.

---

**Supplemental Figure 1.**
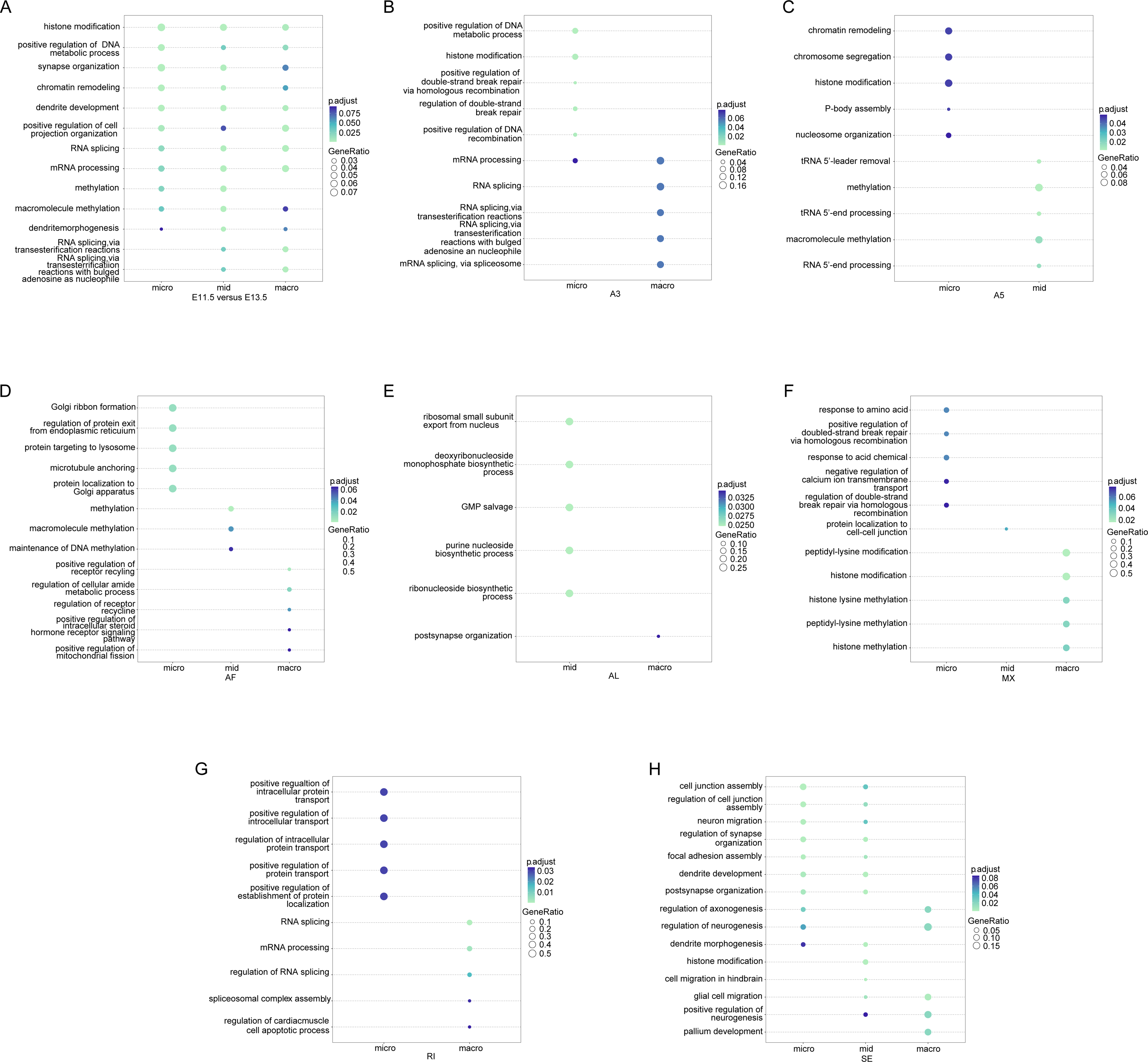
Functional enrichment analysis of differential AS events at gene-level before and after clustering. **A.** Enriched GO terms for three groups of differential AS genes identified between embryonic mouse forebrain E11.5 and E13.5, which are clustered on the size of spliced exons **B-H.** GO enrichment analysis for clustered differential AS genes identified between embryonic mouse forebrain E11.5 and E13.5 after classification into seven types of AS: A3 (B), A5 (C), AF (D), AL (E), MX (F), RI (G), SE (H) GeneRatio: the number of enriched genes over the total number of input genes

**Supplemental Figure 2.**
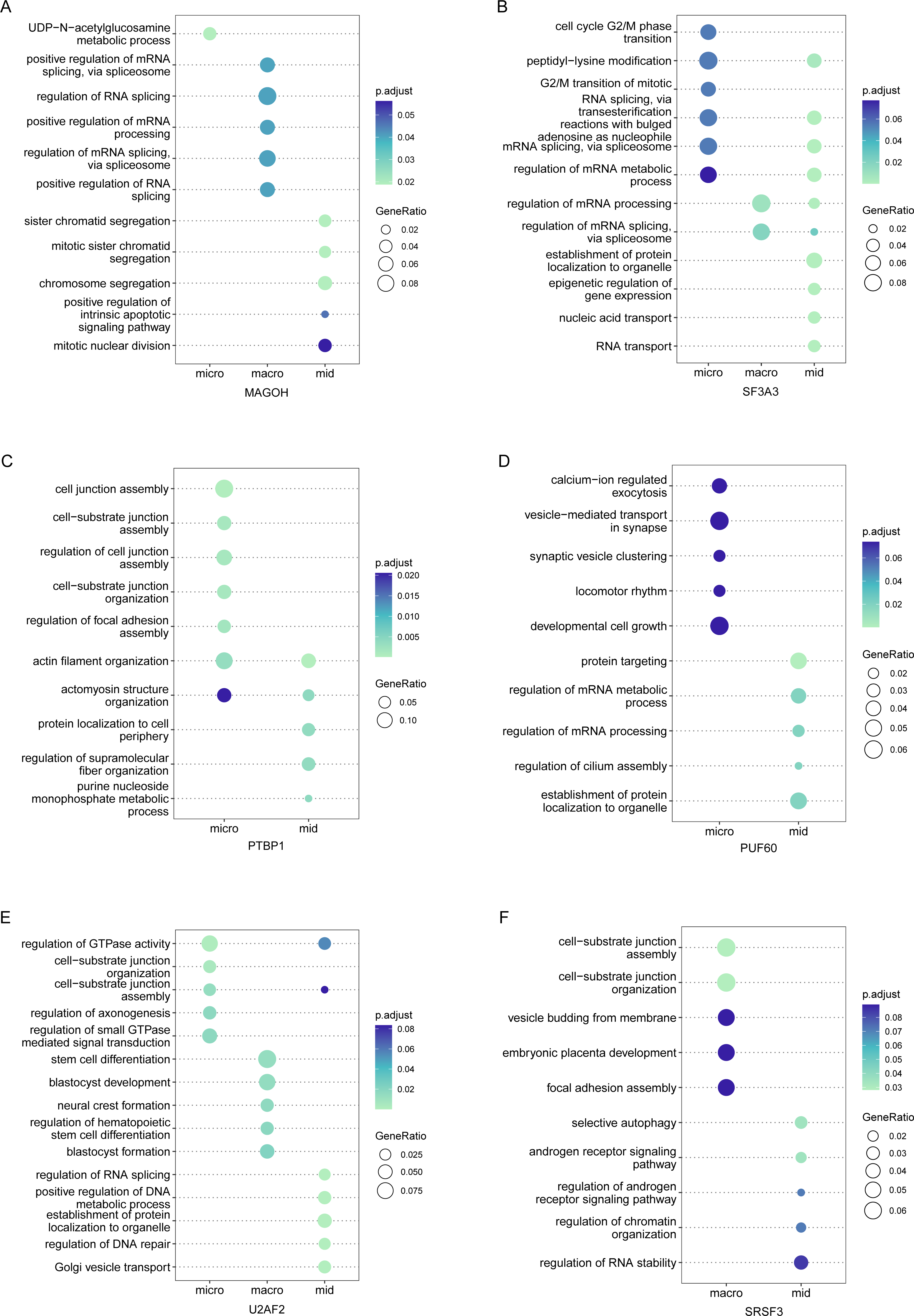
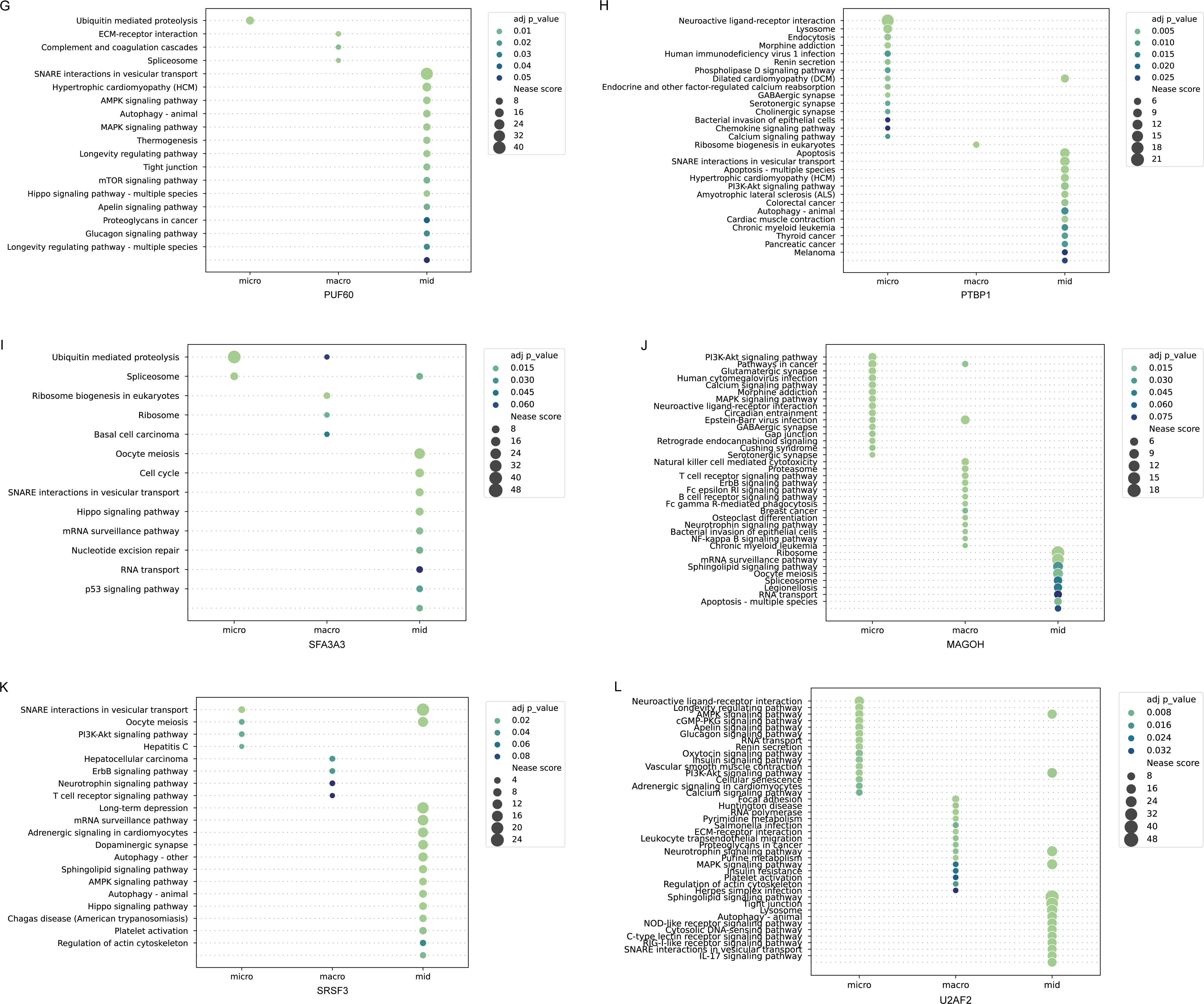
Functional enrichment analysis of differential AS events at gene- and exon-level for the SE type in RBP knockdowns. **A-F.** Enriched GO terms for three groups of differential AS genes in SE for six RBP knockdowns, which are clustered on the size of spliced exons **G-L.** Exon-level KEGG enrichment for three groups of spliced exons in SE for six RBP knockdowns GeneRatio: the number of enriched genes over the total number of input genes

**Supplemental Figure 3.**
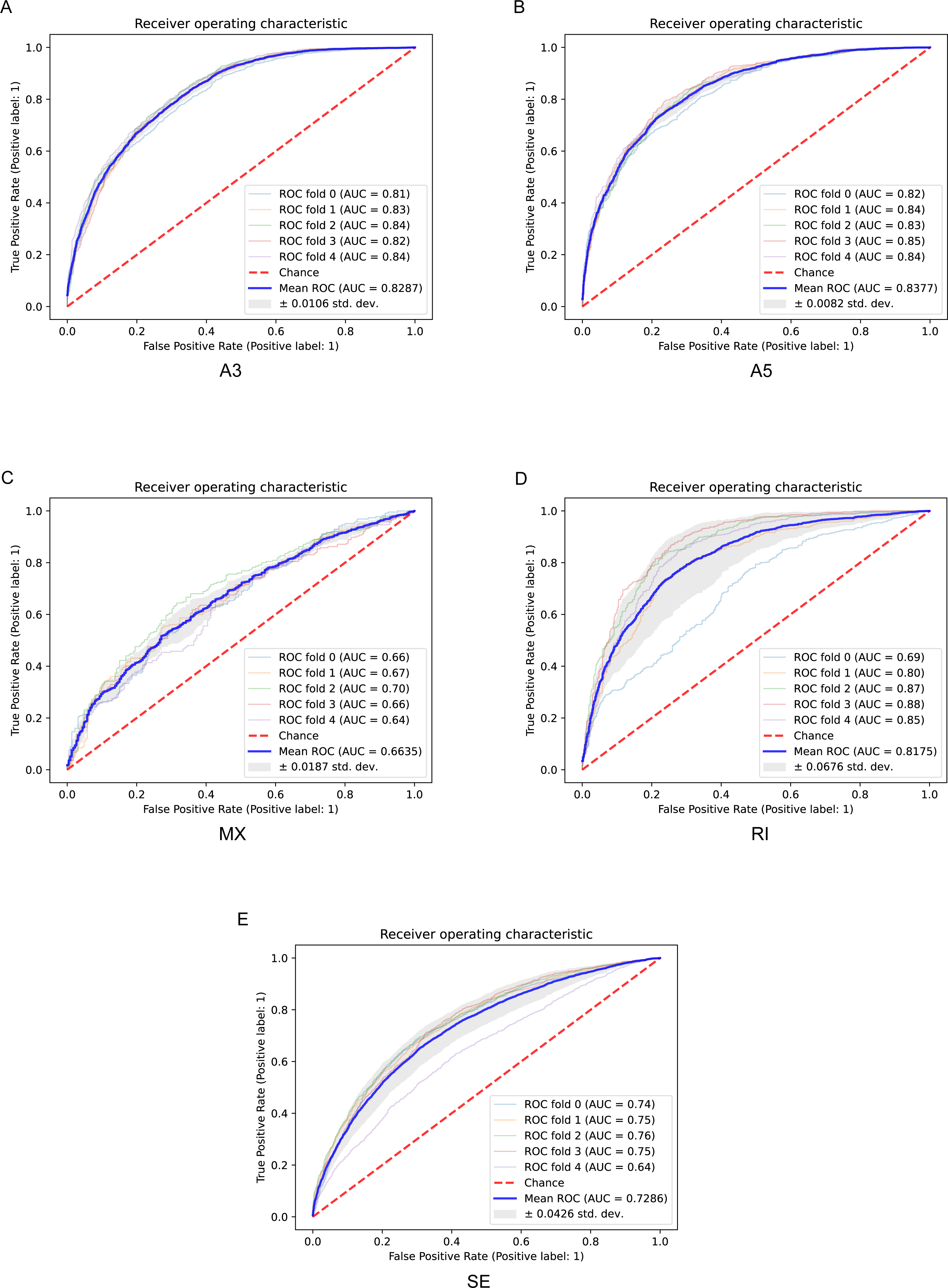
ROC curves of PSI high versus low classifications for five AS types at E16.5 using pooled sequence and epigenetic features (AS data generated by rMATS) **A-E.** ROC was measured with 5-fold cross-validation for each AS type. The mean ROC AUC scores are reported: 0.83 (A3), 0.84 (A5), 0.66 (MX), 0.82 (RI), and 0.73 (SE).

**Supplemental Figure 4.**
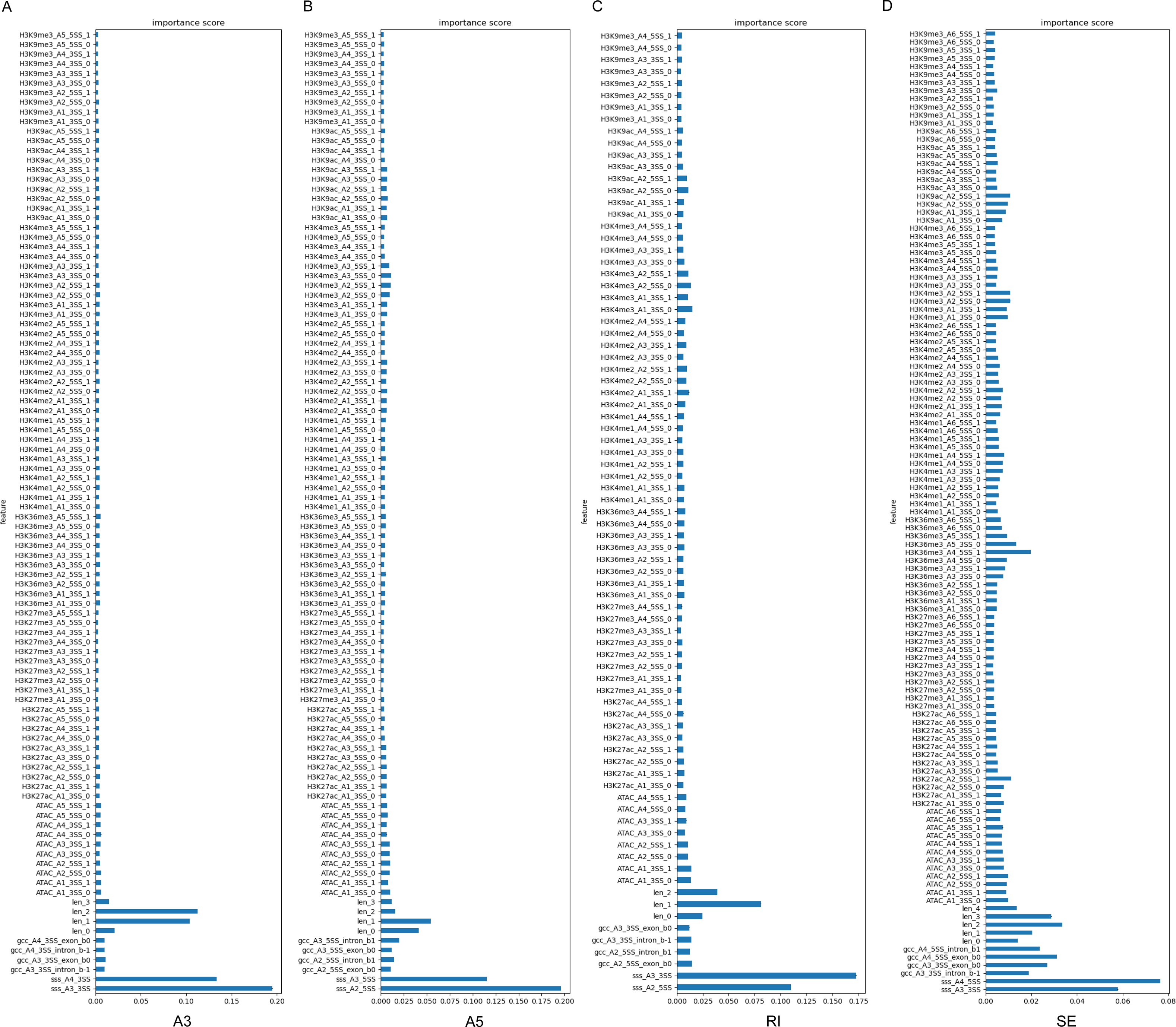
Feature importance in A3, A5, RI, and SE (AS data generated by rMATS) **A-C.** In A3, A5, and RI, the sequence features are most critical in deciding PSI high versus low classification. **D.** In SE, multiple sequence features and epigenetic factors strongly influence PSI high versus low classification.

**Supplemental Table 1.** Accuracies (ACC) and AUC values of random forest models to predict PSI high (top 25%) versus low groups (bottom 25%) for five AS types at each developmental time point using combined sequence features as input (**AS data generated by rMATS**)

|  | E12.5 |  | E13.5 |  | E14.5 |  | E15.5 |  | E16.5 |  |
| --- | --- | --- | --- | --- | --- | --- | --- | --- | --- | --- |
|  | ACC | AUC | ACC | AUC | ACC | AUC | ACC | AUC | ACC | AUC |
| A3 | 0.7454 | 0.8357 | 0.7439 | 0.8339 | 0.7396 | 0.8320 | 0.7432 | 0.8346 | 0.7393 | 0.8287 |
| A5 | 0.7571 | 0.8369 | 0.7641 | 0.8420 | 0.7524 | 0.8319 | 0.7656 | 0.8459 | 0.7571 | 0.8377 |
| MX | 0.6331 | 0.6874 | 0.6197 | 0.6685 | 0.6103 | 0.6685 | 0.5994 | 0.6478 | 0.6169 | 0.6635 |
| RI | 0.7464 | 0.8207 | 0.7513 | 0.8299 | 0.7555 | 0.8303 | 0.7463 | 0.8207 | 0.7413 | 0.8175 |
| SE | 0.6789 | 0.7457 | 0.6828 | 0.7468 | 0.6823 | 0.7465 | 0.6823 | 0.7487 | 0.6687 | 0.7286 |

**Supplemental Table 2.** Accuracies (ACC) and AUC values of random forest models to predict PSI high (top 25%) versus low groups (bottom 25%) for five AS types at E16.5 using each or combined sequence and epigenetic features as input (**AS data generated by rMATS**)

|  | A3 |  | A5 |  | MX |  | RI |  | SE |  |
| --- | --- | --- | --- | --- | --- | --- | --- | --- | --- | --- |
|  | ACC | AUC | ACC | AUC | ACC | AUC | ACC | AUC | ACC | AUC |
| sss | 0.7178 | 0.7819 | 0.7337 | 0.8038 | 0.6093 | 0.6573 | 0.7182 | 0.7792 | 0.5929 | 0.6261 |
| gcc | 0.5413 | 0.5593 | 0.5859 | 0.6158 | 0.5609 | 0.5827 | 0.5606 | 0.5834 | 0.5754 | 0.6060 |
| len | 0.6918 | 0.7623 | 0.6243 | 0.6781 | 0.5689 | 0.5879 | 0.6521 | 0.7021 | 0.5865 | 0.6206 |
| ATAC | 0.5126 | 0.5198 | 0.5517 | 0.5760 | 0.5556 | 0.5665 | 0.5795 | 0.5994 | 0.5693 | 0.5975 |
| H3K4me1 | 0.5076 | 0.5154 | 0.5294 | 0.5423 | 0.5249 | 0.5370 | 0.5781 | 0.6019 | 0.5661 | 0.5939 |
| H3K4me2 | 0.5042 | 0.5137 | 0.5428 | 0.5538 | 0.5471 | 0.5631 | 0.5847 | 0.6060 | 0.5661 | 0.5893 |
| H3K4me3 | 0.5234 | 0.5327 | 0.5464 | 0.5670 | 0.5480 | 0.5663 | 0.5795 | 0.5860 | 0.5685 | 0.5959 |
| H3K9ac | 0.5123 | 0.5211 | 0.5521 | 0.5689 | 0.5369 | 0.5631 | 0.5770 | 0.5917 | 0.5731 | 0.5951 |
| H3K9me3 | 0.5096 | 0.5160 | 0.5222 | 0.5287 | 0.5298 | 0.5333 | 0.5172 | 0.5338 | 0.5420 | 0.5574 |
| H3K27ac | 0.5141 | 0.5182 | 0.5479 | 0.5591 | 0.5244 | 0.5481 | 0.5612 | 0.5792 | 0.5654 | 0.5889 |
| H3K27me3 | 0.5219 | 0.5229 | 0.5157 | 0.5276 | 0.5600 | 0.5683 | 0.5411 | 0.5452 | 0.5384 | 0.5527 |
| H3K36me3 | 0.5220 | 0.5315 | 0.5457 | 0.5654 | 0.5418 | 0.5595 | 0.5612 | 0.5893 | 0.5798 | 0.6143 |
| seq | 0.7676 | 0.8562 | 0.7783 | 0.8612 | 0.6409 | 0.6949 | 0.7538 | 0.8271 | 0.6476 | 0.7019 |
| 8hm+ATAC | 0.5203 | 0.5254 | 0.5574 | 0.5864 | 0.5467 | 0.5677 | 0.5897 | 0.6191 | 0.5994 | 0.6525 |
| 8hm+ATAC+seq | 0.7393 | 0.8287 | 0.7571 | 0.8377 | 0.6169 | 0.6635 | 0.7413 | 0.8175 | 0.6687 | 0.7286 |

sss: splice site strength; gcc: GC content; len: lengths of exons and introns; seq: combined sequence features; hm: histone modifications

**Supplemental Table 3.**
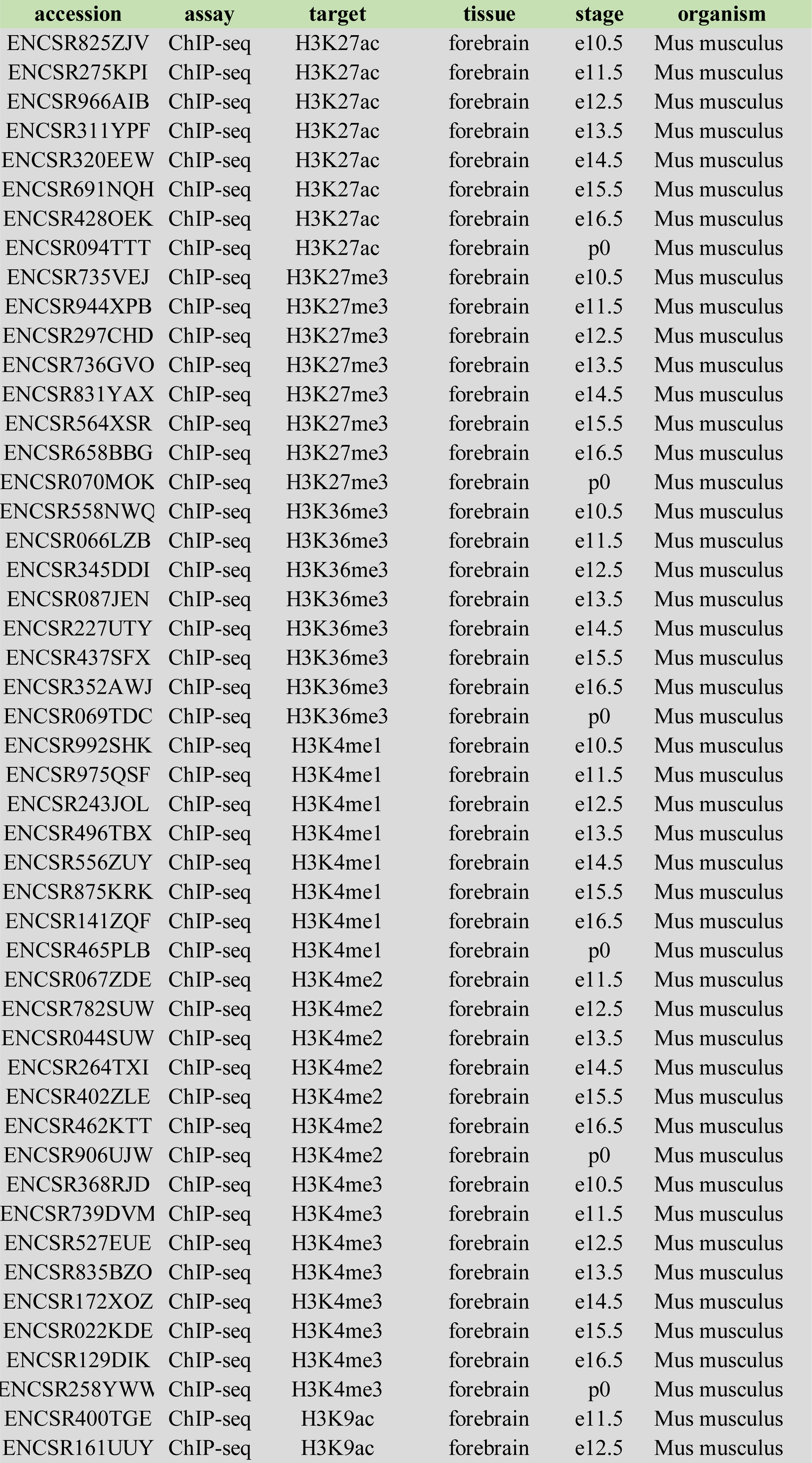
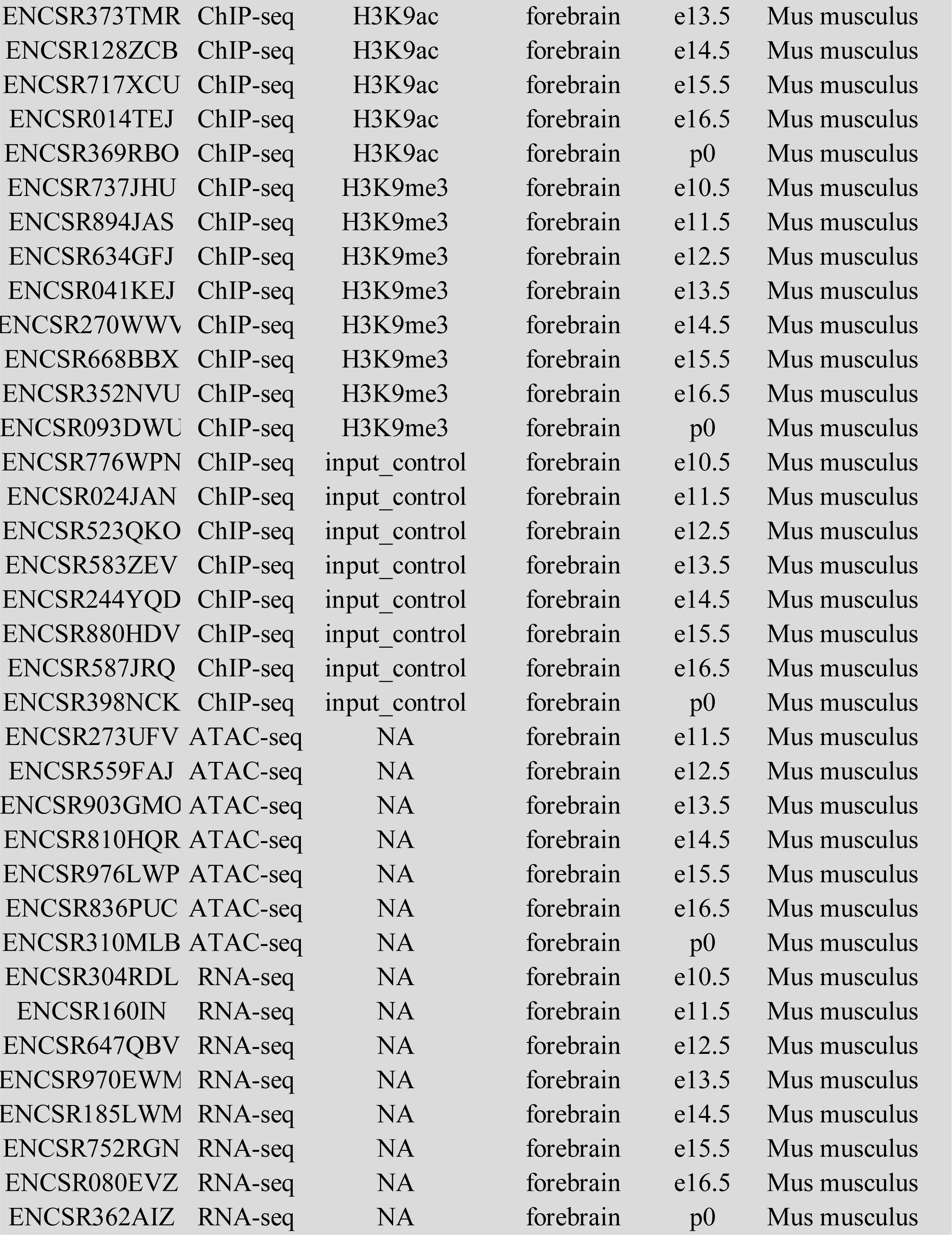
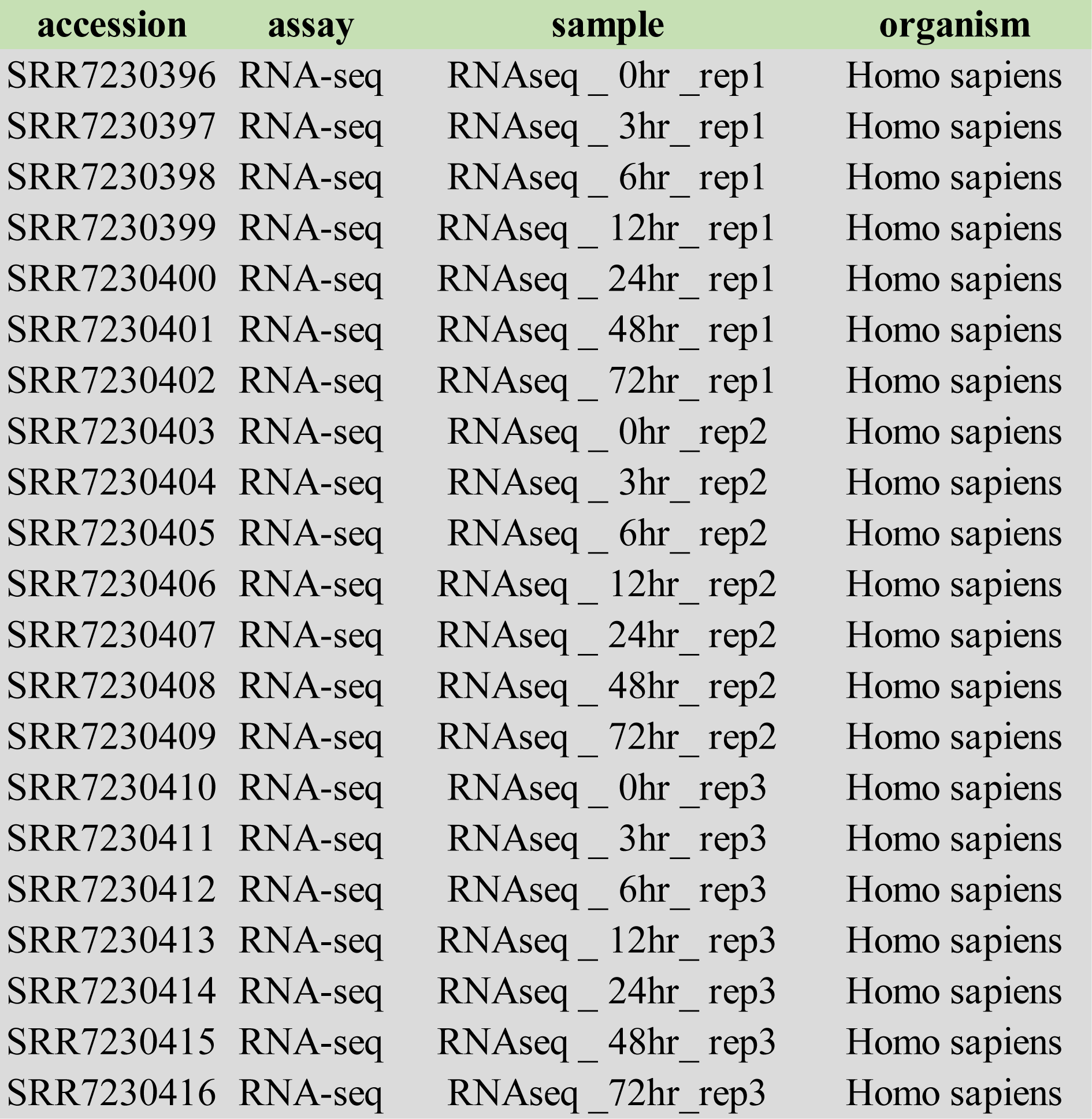
List of accession numbers of RNAseq, ChIPseq, and ATACseq datasets used in our study.

| accession | assay | target | tissue | stage | organism |
| --- | --- | --- | --- | --- | --- |
| ENCSR825ZJV | ChIP-seq | H3K27ac | forebrain | e10.5 | Mus musculus |
| ENCSR275KPI | ChIP-seq | H3K27ac | forebrain | e11.5 | Mus musculus |
| ENCSR966AIB | ChIP-seq | H3K27ac | forebrain | e12.5 | Mus musculus |
| ENCSR311YPF | ChIP-seq | H3K27ac | forebrain | e13.5 | Mus musculus |
| ENCSR320EEW | ChIP-seq | H3K27ac | forebrain | e14.5 | Mus musculus |
| ENCSR691NQH | ChIP-seq | H3K27ac | forebrain | e15.5 | Mus musculus |
| ENCSR428OEK | ChIP-seq | H3K27ac | forebrain | e16.5 | Mus musculus |
| ENCSR094TTT | ChIP-seq | H3K27ac | forebrain | p0 | Mus musculus |
| ENCSR735VEJ | ChIP-seq | H3K27me3 | forebrain | e10.5 | Mus musculus |
| ENCSR944XPB | ChIP-seq | H3K27me3 | forebrain | e11.5 | Mus musculus |
| ENCSR297CHD | ChIP-seq | H3K27me3 | forebrain | e12.5 | Mus musculus |
| ENCSR736GVO | ChIP-seq | H3K27me3 | forebrain | e13.5 | Mus musculus |
| ENCSR831YAX | ChIP-seq | H3K27me3 | forebrain | e14.5 | Mus musculus |
| ENCSR564XSR | ChIP-seq | H3K27me3 | forebrain | e15.5 | Mus musculus |
| ENCSR658BBG | ChIP-seq | H3K27me3 | forebrain | e16.5 | Mus musculus |
| ENCSR070MOK | ChIP-seq | H3K27me3 | forebrain | p0 | Mus musculus |
| ENCSR558NWQ | ChIP-seq | H3K36me3 | forebrain | e10.5 | Mus musculus |
| ENCSR066LZB | ChIP-seq | H3K36me3 | forebrain | e11.5 | Mus musculus |
| ENCSR345DDI | ChIP-seq | H3K36me3 | forebrain | e12.5 | Mus musculus |
| ENCSR087JEN | ChIP-seq | H3K36me3 | forebrain | e13.5 | Mus musculus |
| ENCSR227UTY | ChIP-seq | H3K36me3 | forebrain | e14.5 | Mus musculus |
| ENCSR437SFX | ChIP-seq | H3K36me3 | forebrain | e15.5 | Mus musculus |
| ENCSR352AWJ | ChIP-seq | H3K36me3 | forebrain | e16.5 | Mus musculus |
| ENCSR069TDC | ChIP-seq | H3K36me3 | forebrain | p0 | Mus musculus |
| ENCSR992SHK | ChIP-seq | H3K4me1 | forebrain | e10.5 | Mus musculus |
| ENCSR975QSF | ChIP-seq | H3K4me1 | forebrain | e11.5 | Mus musculus |
| ENCSR243JOL | ChIP-seq | H3K4me1 | forebrain | e12.5 | Mus musculus |
| ENCSR496TBX | ChIP-seq | H3K4me1 | forebrain | e13.5 | Mus musculus |
| ENCSR556ZUY | ChIP-seq | H3K4me1 | forebrain | e14.5 | Mus musculus |
| ENCSR875KRK | ChIP-seq | H3K4me1 | forebrain | e15.5 | Mus musculus |
| ENCSR141ZQF | ChIP-seq | H3K4me1 | forebrain | e16.5 | Mus musculus |
| ENCSR465PLB | ChIP-seq | H3K4me1 | forebrain | p0 | Mus musculus |
| ENCSR067ZDE | ChIP-seq | H3K4me2 | forebrain | e11.5 | Mus musculus |
| ENCSR782SUW | ChIP-seq | H3K4me2 | forebrain | e12.5 | Mus musculus |
| ENCSR044SUW | ChIP-seq | H3K4me2 | forebrain | e13.5 | Mus musculus |
| ENCSR264TXI | ChIP-seq | H3K4me2 | forebrain | e14.5 | Mus musculus |
| ENCSR402ZLE | ChIP-seq | H3K4me2 | forebrain | e15.5 | Mus musculus |
| ENCSR462KTT | ChIP-seq | H3K4me2 | forebrain | e16.5 | Mus musculus |
| ENCSR906UJW | ChIP-seq | H3K4me2 | forebrain | p0 | Mus musculus |
| ENCSR368RJD | ChIP-seq | H3K4me3 | forebrain | e10.5 | Mus musculus |
| ENCSR739DVM | ChIP-seq | H3K4me3 | forebrain | e11.5 | Mus musculus |
| ENCSR527EUE | ChIP-seq | H3K4me3 | forebrain | e12.5 | Mus musculus |
| ENCSR835BZO | ChIP-seq | H3K4me3 | forebrain | e13.5 | Mus musculus |
| ENCSR172XOZ | ChIP-seq | H3K4me3 | forebrain | e14.5 | Mus musculus |
| ENCSR022KDE | ChIP-seq | H3K4me3 | forebrain | e15.5 | Mus musculus |
| ENCSR129DIK | ChIP-seq | H3K4me3 | forebrain | e16.5 | Mus musculus |
| ENCSR258YWW | ChIP-seq | H3K4me3 | forebrain | p0 | Mus musculus |
| ENCSR400TGE | ChIP-seq | H3K9ac | forebrain | e11.5 | Mus musculus |
| ENCSR161UUY | ChIP-seq | H3K9ac | forebrain | e12.5 | Mus musculus |
| ENCSR373TMR | ChIP-seq | H3K9ac | forebrain | e13.5 | Mus musculus |
| ENCSR128ZCB | ChIP-seq | H3K9ac | forebrain | e14.5 | Mus musculus |
| ENCSR717XCU | ChIP-seq | H3K9ac | forebrain | e15.5 | Mus musculus |
| ENCSR014TEJ | ChIP-seq | H3K9ac | forebrain | e16.5 | Mus musculus |
| ENCSR369RBO | ChIP-seq | H3K9ac | forebrain | p0 | Mus musculus |
| ENCSR737JHU | ChIP-seq | H3K9me3 | forebrain | e10.5 | Mus musculus |
| ENCSR894JAS | ChIP-seq | H3K9me3 | forebrain | e11.5 | Mus musculus |
| ENCSR634GFJ | ChIP-seq | H3K9me3 | forebrain | e12.5 | Mus musculus |
| ENCSR041KEJ | ChIP-seq | H3K9me3 | forebrain | e13.5 | Mus musculus |
| ENCSR270WWV | ChIP-seq | H3K9me3 | forebrain | e14.5 | Mus musculus |
| ENCSR668BBX | ChIP-seq | H3K9me3 | forebrain | e15.5 | Mus musculus |
| ENCSR352NVU | ChIP-seq | H3K9me3 | forebrain | e16.5 | Mus musculus |
| ENCSR093DWU | ChIP-seq | H3K9me3 | forebrain | p0 | Mus musculus |
| ENCSR776WPN | ChIP-seq | input_control | forebrain | e10.5 | Mus musculus |
| ENCSR024JAN | ChIP-seq | input_control | forebrain | e11.5 | Mus musculus |
| ENCSR523QKO | ChIP-seq | input_control | forebrain | e12.5 | Mus musculus |
| ENCSR583ZEV | ChIP-seq | input_control | forebrain | e13.5 | Mus musculus |
| ENCSR244YQD | ChIP-seq | input_control | forebrain | e14.5 | Mus musculus |
| ENCSR880HDV | ChIP-seq | input_control | forebrain | e15.5 | Mus musculus |
| ENCSR587JRQ | ChIP-seq | input_control | forebrain | e16.5 | Mus musculus |
| ENCSR398NCK | ChIP-seq | input_control | forebrain | p0 | Mus musculus |
| ENCSR273UFV | ATAC-seq | NA | forebrain | e11.5 | Mus musculus |
| ENCSR559FAJ | ATAC-seq | NA | forebrain | e12.5 | Mus musculus |
| ENCSR903GMO | ATAC-seq | NA | forebrain | e13.5 | Mus musculus |
| ENCSR810HQR | ATAC-seq | NA | forebrain | e14.5 | Mus musculus |
| ENCSR976LWP | ATAC-seq | NA | forebrain | e15.5 | Mus musculus |
| ENCSR836PUC | ATAC-seq | NA | forebrain | e16.5 | Mus musculus |
| ENCSR310MLB | ATAC-seq | NA | forebrain | p0 | Mus musculus |
| ENCSR304RDL | RNA-seq | NA | forebrain | e10.5 | Mus musculus |
| ENCSR160IN | RNA-seq | NA | forebrain | e11.5 | Mus musculus |
| ENCSR647QBV | RNA-seq | NA | forebrain | e12.5 | Mus musculus |
| ENCSR970EWM | RNA-seq | NA | forebrain | e13.5 | Mus musculus |
| ENCSR185LWM | RNA-seq | NA | forebrain | e14.5 | Mus musculus |
| ENCSR752RGN | RNA-seq | NA | forebrain | e15.5 | Mus musculus |
| ENCSR080EVZ | RNA-seq | NA | forebrain | e16.5 | Mus musculus |
| ENCSR362AIZ | RNA-seq | NA | forebrain | p0 | Mus musculus |

| accession | assay | sample | organism |
| --- | --- | --- | --- |
| SRR7230396 | RNA-seq | RNAseq _ 0hr _rep1 | Homo sapiens |
| SRR7230397 | RNA-seq | RNAseq _ 3hr _rep1 | Homo sapiens |
| SRR7230398 | RNA-seq | RNAseq _ 6hr _rep1 | Homo sapiens |
| SRR7230399 | RNA-seq | RNAseq _ 12hr _rep1 | Homo sapiens |
| SRR7230400 | RNA-seq | RNAseq _ 24hr _rep1 | Homo sapiens |
| SRR7230401 | RNA-seq | RNAseq _ 48hr _rep1 | Homo sapiens |
| SRR7230402 | RNA-seq | RNAseq _ 72hr _rep1 | Homo sapiens |
| SRR7230403 | RNA-seq | RNAseq _ 0hr _rep2 | Homo sapiens |
| SRR7230404 | RNA-seq | RNAseq _ 3hr _rep2 | Homo sapiens |
| SRR7230405 | RNA-seq | RNAseq _ 6hr _rep2 | Homo sapiens |
| SRR7230406 | RNA-seq | RNAseq _ 12hr _rep2 | Homo sapiens |
| SRR7230407 | RNA-seq | RNAseq _ 24hr _rep2 | Homo sapiens |
| SRR7230408 | RNA-seq | RNAseq _ 48hr _rep2 | Homo sapiens |
| SRR7230409 | RNA-seq | RNAseq _ 72hr _rep2 | Homo sapiens |
| SRR7230410 | RNA-seq | RNAseq _ 0hr _rep3 | Homo sapiens |
| SRR7230411 | RNA-seq | RNAseq _ 3hr _rep3 | Homo sapiens |
| SRR7230412 | RNA-seq | RNAseq _ 6hr _rep3 | Homo sapiens |
| SRR7230413 | RNA-seq | RNAseq _ 12hr _rep3 | Homo sapiens |
| SRR7230414 | RNA-seq | RNAseq _ 24hr _rep3 | Homo sapiens |
| SRR7230415 | RNA-seq | RNAseq _ 48hr _rep3 | Homo sapiens |
| SRR7230416 | RNA-seq | RNAseq _ 72hr _rep3 | Homo sapiens |

